# β-Amyloid Induces Microglial Expression of GPC4 and APOE Leading to Increased Neuronal Tau Pathology and Toxicity

**DOI:** 10.1101/2025.02.20.637701

**Authors:** Brandon B. Holmes, Thaddeus K. Weigel, Jesseca M. Chung, Sarah K. Kaufman, Brandon I. Apresa, James R. Byrnes, Kaan S. Kumru, Jaime Vaquer-Alicea, Ankit Gupta, Indigo V. L. Rose, Yun Zhang, Alissa L. Nana, Dina Alter, Lea T. Grinberg, Salvatore Spina, Kevin K. Leung, Carlo Condello, Martin Kampmann, William W. Seeley, Jaeda C. Coutinho-Budd, James A. Wells

## Abstract

To elucidate the impact of Aβ pathology on microglia in Alzheimer’s disease pathogenesis, we profiled the microglia surfaceome following treatment with Aβ fibrils. Our findings reveal that Aβ-associated human microglia upregulate Glypican 4 (GPC4), a GPI-anchored heparan sulfate proteoglycan (HSPG). In a *Drosophila* amyloidosis model, glial GPC4 expression exacerbates motor deficits and reduces lifespan, indicating that glial GPC4 contributes to a toxic cellular program during neurodegeneration. In cell culture, GPC4 enhances microglia phagocytosis of tau aggregates, and shed GPC4 can act *in trans* to facilitate tau aggregate uptake and seeding in neurons. Additionally, our data demonstrate that GPC4-mediated effects are amplified in the presence of APOE. These studies offer a mechanistic framework linking Aβ and tau pathology through microglial HSPGs and APOE.

## INTRODUCTION

The defining pathological features of Alzheimer’s disease are the accumulation of β-amyloid (Aβ) plaques and tau neurofibrillary tangles.^1,2^ Rodent and human studies suggest that Aβ accelerates the propagation of tau pathology within brain networks, likely through both local and remote amyloid-tau interactions^3–8^. Indeed, the anti-Aβ monoclonal antibodies, lecanemab and donanemab, reduce the deposition of tau pathology in patients with Alzheimer’s disease (AD), possibly via the removal of upstream amyloid plaques.^9–11^ However, the cellular and molecular mechanisms by which Aβ incites the spread of tau pathology remain unknown.

As the resident immune cells of the brain, microglia are the frontline reporters and executioners of inflammatory activity and injury response in neurodegeneration. Emerging evidence now links microglia biology to various neurodegenerative disorders, including AD.^13^ Multi-omics analyses – including GWAS, gene expression data, epigenetic annotations, cellular pathway analyses, and cell-type specific promoter mapping – show that many common AD risk variants are either expressed uniquely in microglia or enriched in microglia-specific enhancer elements.^12–15^ Further, integrative network analyses demonstrate that AD genetic variants converge on microglia biological processes such as phagocytosis, endolysosomal processing, and lipid metabolism.^16,17^ Thus, a substantial body of genetic data implicate a central role for microglia in the pathogenesis of AD.

Emerging experimental data suggests that reactive microglia contribute to the deposition of pathological tau as well as tau-mediated neurodegeneration in AD^18–21^, possibly via APOE.^22,23^ Human PET imaging studies demonstrate that tau propagation pathways depend on microglia networks, and the co-occurrence of Aβ, tau, and microglia reactivity is a stronger predictor of cognitive impairment than the combination of Aβ and tau alone.^24^ Given that microglia reactivity may precede tau pathology^25,26^, we hypothesize that AD reactive microglia are *primed* to accelerate pathological tau spread and that this priming may result from Aβ pathology.

To interrogate how microglia respond to Aβ, we profiled the iPSC-derived microglia surfaceome with surface biotinylation using WGA-HRP glycoproteomics^27^ after treatment with recombinant Aβ fibrils. We found that Aβ stimulates microglia to remodel their cell-surface and specifically express Glypican 4 (GPC4), a GPI-anchored heparan sulfate proteoglycan not previously implicated in microglia biology. In AD patients, microglia associated with Aβ plaques upregulate GPC4, and *in vivo*, glial GPC4 expression accelerates amyloid-related toxicity. The expression of GPC4 enhances tau aggregate uptake in microglia and neurons, and, in concert with APOE, potentiates intraneuronal tau seeding and pathology. These findings motivate the development of novel microglia-targeted therapeutic interventions to disrupt Aβ-induced toxicity and spread of tau pathology.

## RESULTS

### iTF Microglia surfaceomics reveals Aβ fibril-induced upregulation of heparan sulfate proteoglycans and glypicans

The pathology of AD elicits a profound microglial reaction, generating unique transcriptional and functional microglial features.^28,29^ Since microglia are highly dynamic cells capable of altering their receptor profiles in response to tissue injury^30,31^, we aimed to investigate Aβ-associated surfaceome remodeling. We performed cell-surface proteomics on human iPSC-derived induced-transcription factor microglia (iTF Microglia).^32^ The iTF microglia recapitulate core microglia biology, as assessed by the expression of microglia markers and functional responses to lipopolysaccharide (LPS) and phagocytic substrates (**Figures S1A – E**). Given that microglia rapidly react to their microenvironment with altered protein expression, we employed an efficient surfaceome labeling strategy previously developed by our lab.^27^ This technology uses horseradish-peroxidase tethered to wheat-germ agglutinin (WGA-HRP) to bind the cell-surface and biotinylate resident membrane proteins.

To elucidate AD-associated surfaceome changes, we treated iTF Microglia with recombinant Aβ_40_ fibrils, Aβ_42_ fibrils (**Figure S1F**) or lipopolysaccharide (LPS) for 72 h to allow surfaceome remodeling. LPS was used to elicit a generic innate immune response, providing a comparison to amyloid-specific proteomic changes.

Compared to vehicle-treated iTF Microglia, Aβ_40_ fibrils resulted in the significant upregulation (> 1.5-fold) of 30 proteins and downregulation of 62 proteins (**Figure 1A**); Aβ_42_ fibrils led to the significant upregulation of 28 proteins and downregulation of 27 proteins (**Figure 1B**). Of the differentially expressed proteins, 13 (9.7%) were shared between Aβ_40_ and Aβ_42_ fibril-treated iTF Microglia. LPS led to the upregulation of 46 proteins and downregulation of 47 proteins (**Figure 1C**). Only 8 (5.42%) and 11 (8.03%) of these differentially expressed proteins were shared between Aβ_40_ and Aβ_42_ fibril-treated iTF Microglia, respectively; and only 3 (2.8%) proteins were shared between all three treatments. To evaluate for thematic surfaceome changes, we employed Gene Set Enrichment Analysis (**Figure 1D**). Among the top 25 most statistically significant Gene Ontology terms, five implicated a role in proteoglycan biology, including glycosaminoglycan binding (GO:0005539), aminoglycan biosynthetic process (GO:0006023), aminoglycan catabolic process (GO:0006027), aminoglycan metabolic process (GO:0006022), and heparin binding (GO:0008201). Within the proteins comprising these groups of Gene Ontology terms, our mass spectrometry identified 8 proteoglycans on iTF Microglia, six of which were cell-surface bound (**Figure 1E**), and four of which were in the glypican family of heparan sulfate proteoglycans (HSPGs). These glypican HSPGs, particularly Glypican 4 (GPC4), were enriched in the Aβ_40_ and Aβ_42_ fibril-treated iTF Microglia but not in the LPS-treated cells suggesting that their expression may be specifically induced by amyloid pathology. A role for glypicans in microglia has not been previously defined in health or disease.

**Figure 1:**
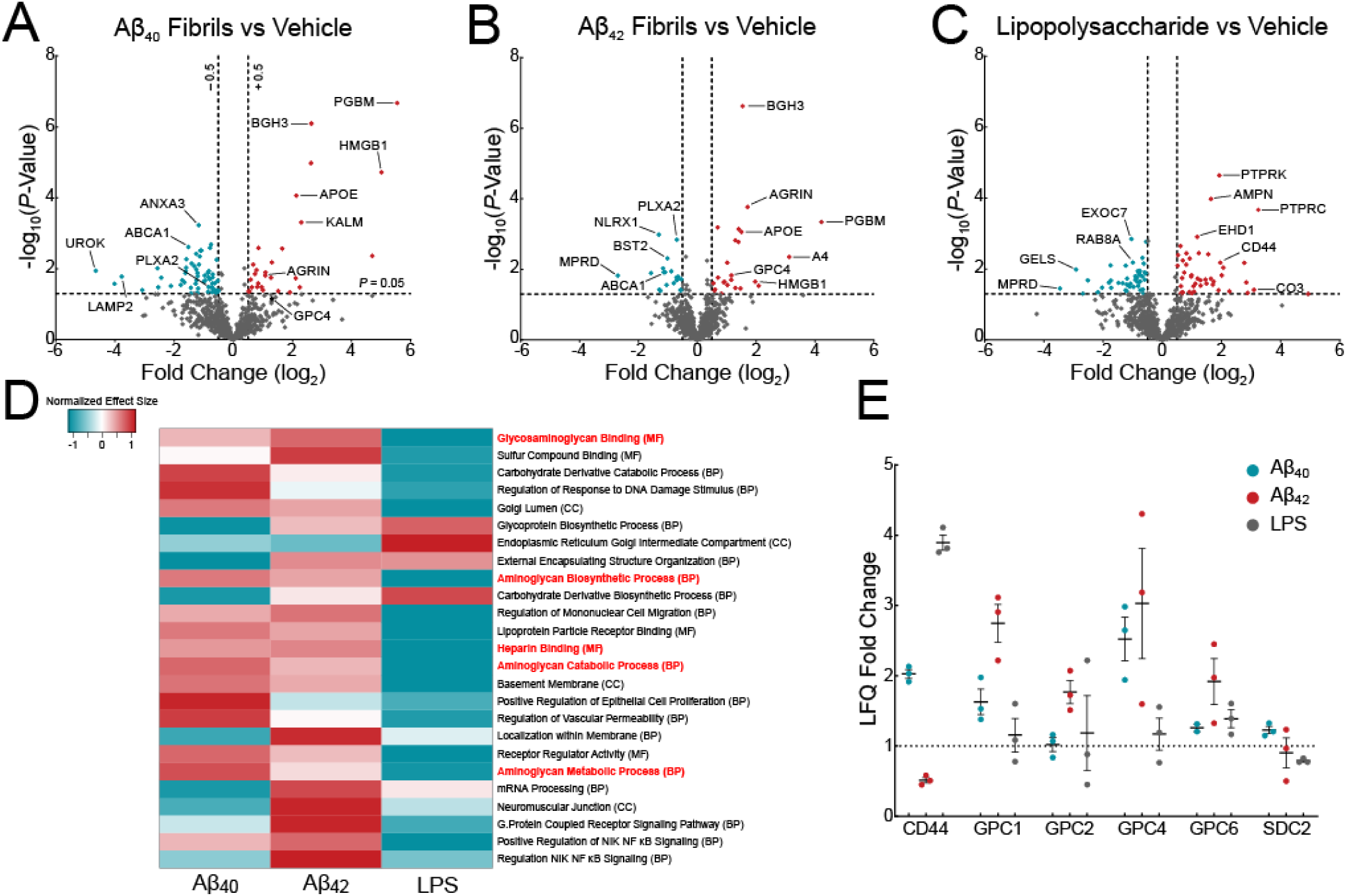
Surfaceomics of iTF microglia reveal upregulation of glypicans. Volcano plots showing changes in surface proteins from iTF Microglia treated with (A) 1 µM Aβ_40_ fibrils, (B) 1 µM Aβ_42_ fibrils, or (C) 100 ng/mL LPS. The - log_10_ transformed p-value is plotted against the log_2_ transformed label-free quantitation ratios (log_2_ fold changes). N = 3 - 6 biological replicates. (D) Gene Set Enrichment Analysis of iTF Microglia treated with Aβ_40_ fibrils, Aβ_42_ fibrils, or LPS. The heatmap shows the normalized effect size of the top 25 most statistically altered pathways. Positive normalized effect size is shown in red (upregulation), and negative normalized effect size is shown in blue (downregulation). Gene Ontology terms in red font relate to glycosaminoglycan and proteoglycan biology. (E) Mass spectrometry label-free quantitation of cell-surface heparan sulfate proteoglycans found in the surfaceomics experiments (A, B, C).

### iTF Microglia uniquely and specifically upregulate heparan sulfate and GPC4 in response to Aβ fibrils

HSPGs have been previously recognized to co-deposit within extracellular amyloid plaques,^33,34^ as well as play a role in tau protein aggregate uptake and intracellular seeding in neurons.^35–37^ To further explore the involvement of HSPGs and glypicans in AD-related microglia, we directly tested whether Aβ fibrils can result in increased cell-surface heparan sulfate in microglia. iTF microglia treated with Aβ_40_ or Aβ_42_ fibrils for 24 h showed a dose-dependent increase, up to four-fold, in cell-surface heparan sulfate as measured by flow cytometry using an anti-HSPG antibody (clone 10E4) (**Figures 2A and 2B**). Importantly, treatment with Aβ_42_ monomer did not result in increased cell-surface heparan sulfate (**Figure S2A**) suggesting that amyloids, but not cognate monomer, have the capacity to induce cell-surface HSPGs. Furthermore, iTF Microglia treated with innate immune activators TNFα (1 nM), IFNβ (100 pM), IFNγ (100 pM), LPS (100 ng/mL), BioParticles (10 µg/mL), the heparan sulfate-binding TAT peptide (5 µM), or scrambled Aβ_42_ peptide for 24 hours also did not increase cell-surface heparan sulfate (**Figure 2C**) suggesting that generic inflammation is insufficient to increase microglial HSPGs. To determine if microglia increase their heparan sulfate in response to other amyloids, we treated iTF Microglia with tau monomer, synuclein monomer, tau fibrils, or synuclein fibrils (**Figure S1F**) and again quantified heparan sulfate. Both tau fibrils and synuclein fibrils, but not cognate monomers, induced the expression of cell-surface heparan sulfate further confirming that amyloids, but not their monomeric equivalent, are competent to elicit this HSPG response (**Figure S2B – S2E**).

**Figure 2:**
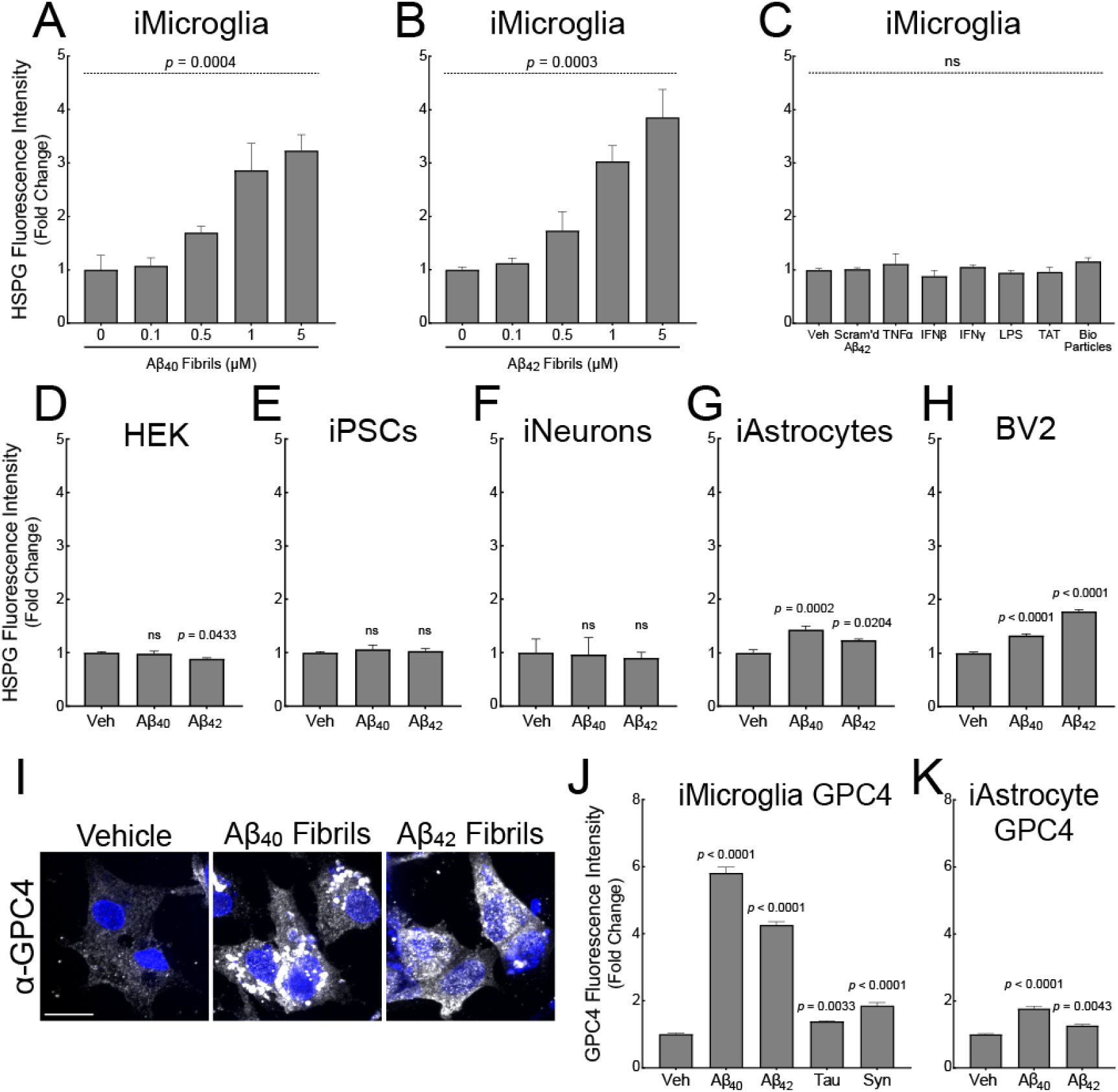
Amyloids induce iTF Microglial cell-surface heparan sulfate and GPC4 expression. iTF-Microglia treated with (A) Aβ_40_ and (B) Aβ_42_ fibrils have dose-dependent increases in cell-surface heparan sulfate as measured by flow cytometry with 10E4 antibody. (C) Other inflammatory substrates and controls do not alter cell-surface heparan sulfate levels. The statistical analyses were performed with a one-way ANOVA. N = 3 biological replicates. Heparan sulfate cell-surface staining with 10E4 on (D) HEK293T cells, (E) iPSCs, (F) iNeurons, (G) iAstrocytes, and (H) BV2 cells treated with 1 µM Aβ_40_ and Aβ_42_ fibrils. The statistical analyses were performed with a one-way ANOVA and Holm-Sidak multiple comparisons tests for the adjusted *p*-values. N = 3 biological replicates. (I) GPC4 immunocytochemistry of iTF Microglia treated with 1 µM Aβ_40_ or Aβ_42_ fibrils. Scale bar = 10 µm. GPC4 flow cytometry quantification of (J) iTF Microglia or (K) iAstrocytes treated with 1 µM proteopathic amyloid fibrils. The statistical analyses were performed with a one-way ANOVA and Holm-Sidak multiple comparisons tests for the adjusted *p*-values. N = 3 biological replicates. In all graphs, the data represent the means ± SEM.

We next investigated whether Aβ fibrils can prime other cell types to upregulate cell-surface heparan sulfate. We treated HEK293T cells, undifferentiated iPSCs (precursors to iTF Microglia), iNeurons, iAstrocytes, and mouse BV2 cells with 1 µM of Aβ_40_ fibrils or Aβ_42_ fibrils for 24 h. HEK293T cells, iPSCs, and iNeurons did not increase cell-surface heparan sulfate by 10E4 staining (**Figure 2D – 2F**). iAstrocytes and BV2 cells responded to Aβ fibrils with a modest (<2-fold) increase in cell-surface heparan sulfate (**Figure 2G – 2H**). This data suggest that human microglia may be particularly responsive to Aβ-induced heparan sulfate expression.

Cell-surface heparan sulfates are known to be attached to glypican (GPC1 – 6) and syndecan (SDC1 – 4) core protein family members. From our iTF Microglia surfaceomics, we identified four out of six glypican family members. GPC4, in particular, was significantly upregulated upon Aβ_42_ fibril treatment. To test if Aβ fibrils directly upregulate cell-surface glypicans, we treated iTF microglia with 1 µM Aβ_40_ or Aβ_42_ fibrils for 24 h. Aβ_40_ and Aβ_42_ fibrils robustly elevated GPC4 expression as measured by microscopy, flow cytometry, and qPCR whereas tau fibrils and synuclein fibrils only modestly elevated GPC4 (**Figure 2I, 2J and S2F**). Further, the same Aβ fibril treatment led to minimal increases in iAstrocyte GPC4 (**Figure 2K**). These data support the concept that microglia, more so than astrocytes, undergo Aβ-induced GPC4 expression. Finally, we evaluated the effect of Aβ fibril treatment on iTF Microglia cell-surface expression for the remainder of the glypican family. GPC2, GPC3, and GPC5 were weakly expressed in iTF Microglia and unchanged after treatment with Aβ fibrils (**Figure S2G**). GPC6 expression was elevated after treatment with Aβ fibrils, albeit to a much smaller degree than GPC4 (**Figure S2G and S2H**). We could not detect the presence of GPC1 (data not shown). Taken together, this data demonstrate that recombinant Aβ fibrils robustly and specifically upregulate microglia GPC4 expression in the iPSC culture system.

### Human microglia GPC4 expression correlates with Aβ plaque pathology in Alzheimer’s disease

To investigate whether microglia express GPC4 in neurodegeneration, we performed immunohistochemistry on human brain sections containing Aβ plaques and neurofibrillary tangles (Alzheimer’s disease), neurofibrillary tangles only (corticobasal degeneration), Aβ plaques only, or neither, (healthy age-matched control tissue) (**Table 1**). We stained human tissue with IBA1 and GPC4 antibodies and measured microglia GPC4 expression. Aged-matched control microglia expressed minimal GPC4 whereas AD microglia robustly expressed GPC4 (**Figure 3A**). Quantification of microglia-specific expression of GPC4 revealed a 2.2-fold increase in AD samples (**Figure 3B**). Additionally, we observed some punctate and low-intensity GPC4 expression in non-microglia cells.

**Figure 3:**
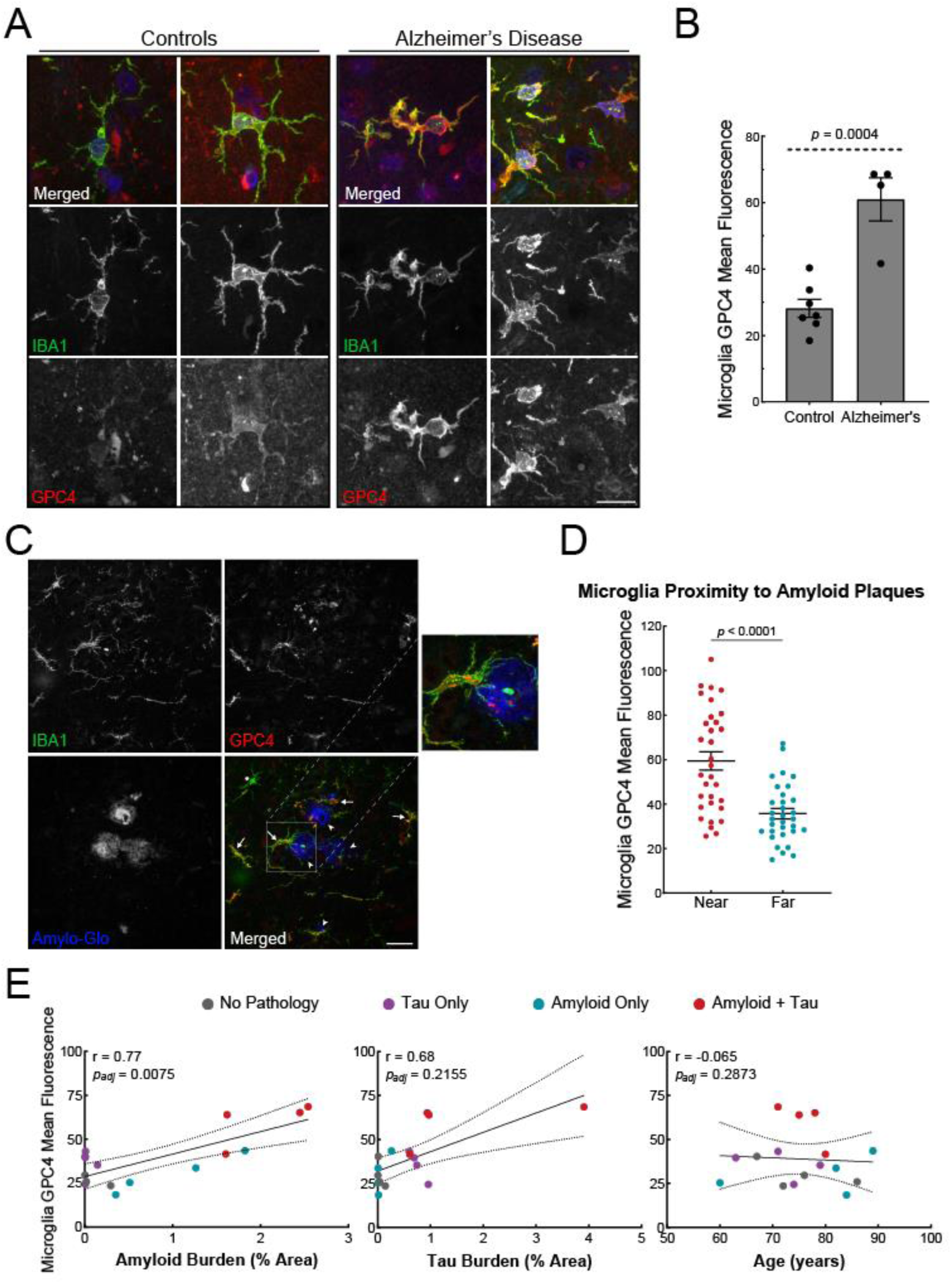
Microglial GPC4 expression is upregulated in human AD brain and correlates with amyloid pathology. (A) Representative confocal images of IBA1 (green), GPC4 (red), and DAPI (blue) in two age-matched controls and two AD cases. Scale bar = 20 µm. (B) GPC4 mean fluorescence intensity measurements in IBA1^+^ microglia from age-matched controls (N = 7) and AD (N = 4) cases. The statistical analysis was performed with a Student t-test for averaged values from individual subjects. The data represent the means ± SEM. (C) Representative confocal images of IBA1 (green), GPC4 (red), and Amylo-Glo (blue) in an AD case. Arrowhead = amyloid plaque; Arrow = GPC4^+^ microglia; Asterisk = GPC4^−^ microglia. Scale bar = 40 µm. (D) Quantification of GPC4 mean intensity values in IBA^+^ microglia located at two distances from Amylo-Glo^+^ Aβ plaques. A total of eight plaques were measured from each brain, and for each plaque, microglia were binned into two separate categories, “near” or “far”, based on proximity to the plaque. Near: < 125 µm from center of plaque; Far: > 150 µm and < 350 µm from edge of plaque. The *p*-values were determined by a paired t-test. The data represent the means ± SEM. (E) Scatter plots of microglial GPC4 mean fluorescence intensity per each patient versus their amyloid plaque burden, neurofibrillary tangle burden, or age. Pathological categories colored grey for control, purple for individuals for tau pathology only, blue for individuals with Aβ pathology only, or red for individuals with both tau and Aβ pathology. The *p*-values were determined by Pearson’s correlation with 95% confidence bands and adjusted for confounding by applying multiple linear regression.

We next investigated whether microglia that are near amyloid plaques express greater levels of GPC4 relative to microglia that are far from plaques. We stained brain sections with IBA1 and GPC4 antibodies as well as Amylo-Glo, a fluorescent dye that labels amyloid plaques (**Figure 3C**).^38^ For each AD brain, we selected 8 Amylo-Glo-positive plaques and binned microglia into two groups: near microglia (< 125 µM from the center of the index plaque) or far microglia (> 150 µm from the edge of all plaques and < 350 µm from edge of the index plaque). Microglia near Aβ plaques had significantly higher GPC4 expression compared to microglia far from (**Figure 3D**). Notably, this trend persisted in human brains containing Aβ plaque pathology without neurofibrillary tangles suggesting that microglia GPC4 expression is independent of tau pathology (**Figure S3A**).

We next assessed whether microglia GPC4 expression correlated with amyloid plaque burden or neurofibrillary tangle burden after staining the tissue with an anti-Aβ antibody (DE2) or anti-phosphorylated tau antibody (CP13). After adjusting the GPC4-amyloid pathology correlation for tau pathology as a confounder by applying multiple linear regression, there was a significant relationship between GPC4 expression and amyloid pathology (r = 0.77; *p* = 0.0075), but not for tau pathology (r = 0.68; *p* = 0.2155). There was no correlation between microglia GPC4 expression and patient age (**Figure 3E**).

Finally, we investigated whether other members of the Glypican family were upregulated in AD microglia. We found that GPC1, GPC2, and GPC6 were globally upregulated in human AD tissue compared to healthy aged-matched control. However, GPC1, GPC2, and GPC6, were not expressed by microglia (**Figure S3B**), but rather expressed in cells with a neuronal morphology. Taken together, our data indicate that human AD microglia express GPC4 which is correlated with and proximal to Aβ pathology.

### Mouse microglia do not upregulate GPC4 in models of amyloidosis

We next investigated whether mouse models of amyloidosis recapitulate this phenotype. We performed immunohistochemistry to examine microglia GPC4 expression within the frontal lobe cortex and dentate gyrus of aged non-transgenic C57BL/6J, 5xFAD, and App-SAA knock-in mice. Surprisingly, none of the three mouse lines examined expressed GPC4 in IBA1^+^ microglia (**Figure S4A – S4E**). However, punctate and intense GPC4 expression was detected in GFAP^+^ astrocytes, but this expression was not significantly different between non-transgenic and transgenic mice. These data demonstrate that mouse models of amyloidosis do not recapitulate human microglia GPC4 expression, but rather display persistent astrocyte expression in adulthood. Recent research reveals significant evolutionary divergence between rodent and primate microglia with human microglia exhibiting the greatest transcriptional heterogeneity and number of cellular states.^39,40^ This implies that rodent microglia may have a distinct reactivity profile in comparison to humans, although current rodent studies have not directly evaluated disease-relevant conditions. Therefore, it is possible that the human upregulation of microglial GPC4 in AD is yet another example of evolutionary divergence between rodents and primates.

### Glial GPC4 enhances toxicity in a Drosophila Model of Amyloidosis

Since microglial GPC4 expression is absent from rodent models of amyloidosis, we employed *Drosophila* to control the glial expression of *Dlp*, the fly homolog of human GPC4. We first investigated whether glial expression of GPC4 modifies amyloid toxicity *in vivo* by using a previously established transgenic *Drosophila* models of amyloidosis^41^ (**Figure 4A and S5A**). The expression of human Aβ_42_ in *Drosophila* neurons led to profound and progressive climbing deficits, a measure of motor dysfunction (**Figure 4B, 4C**), as well as premature lethality (**Figure 4D**). Glial over-expression of *Dlp* progressively worsened climbing at day-one and day-five post-eclosion (**Figure 4B and 4C**) as well as reduced lifespan (**Figure 4D**) beyond what was observed for Aβ_42_ alone. Conversely, knockdown of glial *Dlp* with RNAi improved climbing (**Figure 4B and 4C**) and prolonged lifespan (**Figure 4D**). The effect of *Dlp* was amyloid-specific, since these phenotypes were not observed to the same degree in *Drosophila* that do not express human Aβ_42_ (**Figure S5B – S5D**).

**Figure 4:**
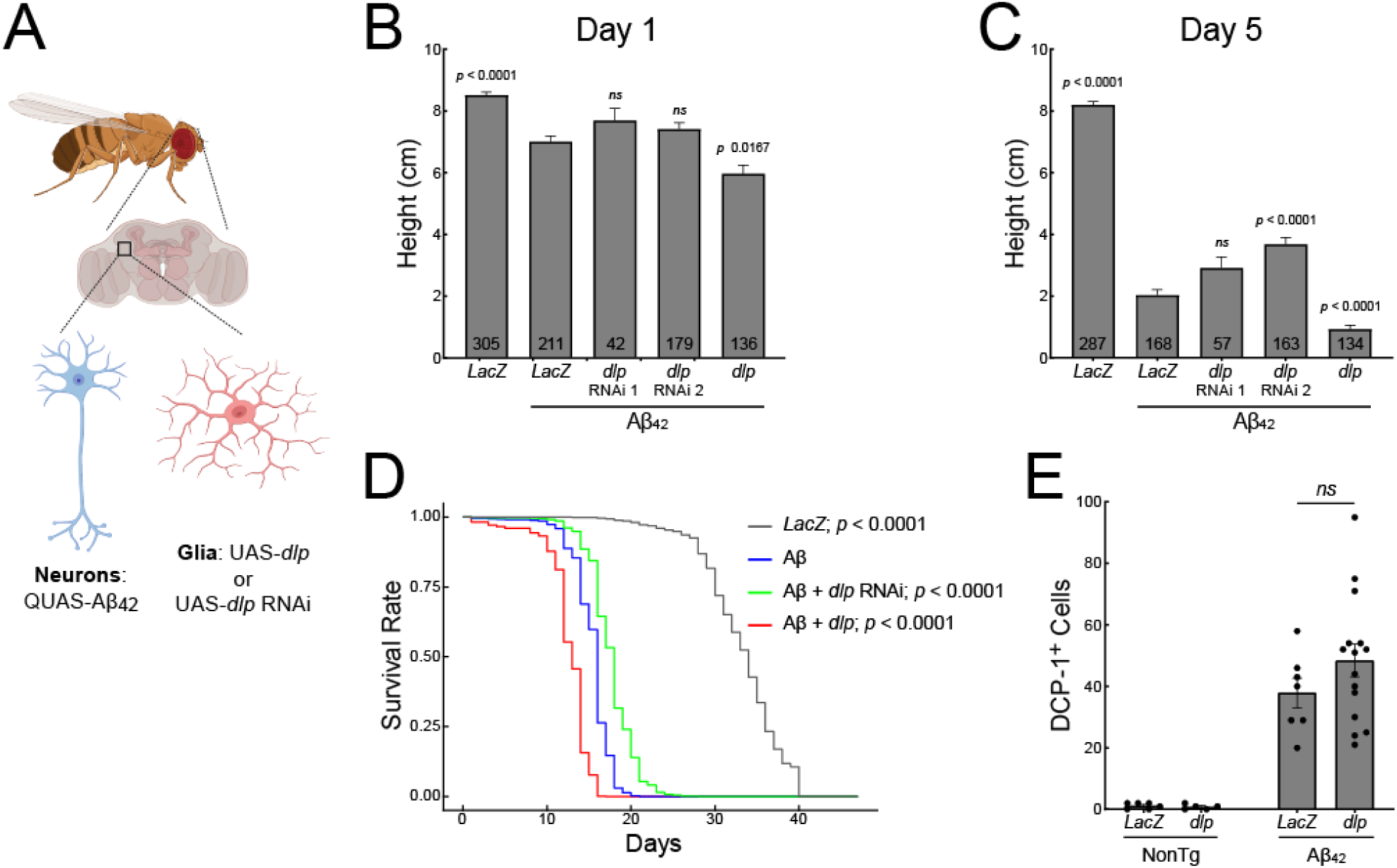
Glial GPC4 worsens climbing and early lethality in an amyloid model of *Drosophila*. (A) Schematic model depicting *Drosophila* transgenic lines in which neurons express Aβ_42_ via the QUAS promoter and glia express *dlp* or *dlp* RNAi via the UAS promoter. Climbing heights were measured at day 1 post-eclosion (B) or day 5 post-eclosion (C) in Aβ_42_ flies expressing LacZ, dlp RNAi, or dlp cDNA. The numbers within the bars represent the number of flies measured per condition. The statistical analyses were performed with a Tobit regression with Bonferroni correction. (D) Median fly lifespans measured in days after eclosion (LacZ = 34 d; Aβ_42_ = 16 d; Aβ_42_ + *dlp* RNAi = 18 d; Aβ_42_ + *dlp* = 12 d). Lifespan data was analyzed using a Cox proportional hazard model with Bonferroni corrections. (LacZ n = 174; Aβ_42_ n = 134; Aβ_42_ + *dlp* RNAi n = 56; Aβ_42_ + *dlp* n = 102). (E) Drosophila brains were immunostained for cleaved Dcp-1 and positive cells were counted across the entire brain. The statistical analysis was performed with a Student t-test. The error bars represent the SEM values.

We next investigated whether the toxic effects of *Dlp* over-expression is mediated via cell death. We stained transgenic *Drosophila* at 10 days post eclosion with an anti-DCP-1 antibody to detect activation of caspase. We did not observe a significant increase in apoptosis in *Drosophila* over-expressing glial *Dlp* (**Figure 4E**) suggesting that the toxic effects of *Dlp* on amyloidosis may be due to mechanisms other than this form of cell death. Taken together, these data demonstrate that glial-derived *Dlp*, the ortholog of human GPC4, enhance Aβ toxicity *in vivo*.

### GPC4 mediates tau phagocytosis in iTF Microglia

Having confirmed that GPC4 is upregulated in human AD-associated microglia and potentiates toxicity *in vivo*, we next investigated the impact of GPC4 on microglial function. Our prior work in neurons demonstrates that HSPGs govern the internalization and propagation of tau pathology.^35,36,42^ However, the specific core proteins mediating this response remain unknown. Here, we hypothesized that Aβ fibril-induced GPC4 expression may facilitate tau aggregate phagocytosis in microglia. To investigate this, we primed iTF microglia with varying concentrations of Aβ_40_ or Aβ_42_ fibrils for 24 h, washed the cells, applied pHrodo-labeled tau fibrils, and monitored phagocytosis over time. We observed a significant increase in tau phagocytosis when iTF microglia were primed with increasing doses of Aβ fibrils (**Figure 5A and 5B**). This Aβ-induced tau phagocytosis was inhibited by ∼50% by the co-application of heparin or chlorate, inhibitors of heparan sulfate proteoglycans (**Figure 5C and 5D**) demonstrating that heparan sulfate contributes to Aβ priming of iTF microglia. Aβ priming did not alter the phagocytosis of magnetic beads (**Figure S6A**) which are not known to enter microglia in an HSPG-dependent manner. To specifically assess if GPC4 mediates tau aggregate phagocytosis, we employed iTF Microglia cells expressing trimethoprim-inducible CRISPRi and CRISPRa machinery to downregulate and upregulate GPC4, respectively. Downregulation of GPC4 via CRISPRi reduced tau fibril phagocytosis, whereas upregulation of GPC4 via CRISPRa increased phagocytosis (**Figure 5E and 5F**). The extent of tau fibril phagocytosis was proportional to the degree of GPC4 expression change as measured by flow cytometry (**Figure S6C and S6D)**.

**Figure 5:**
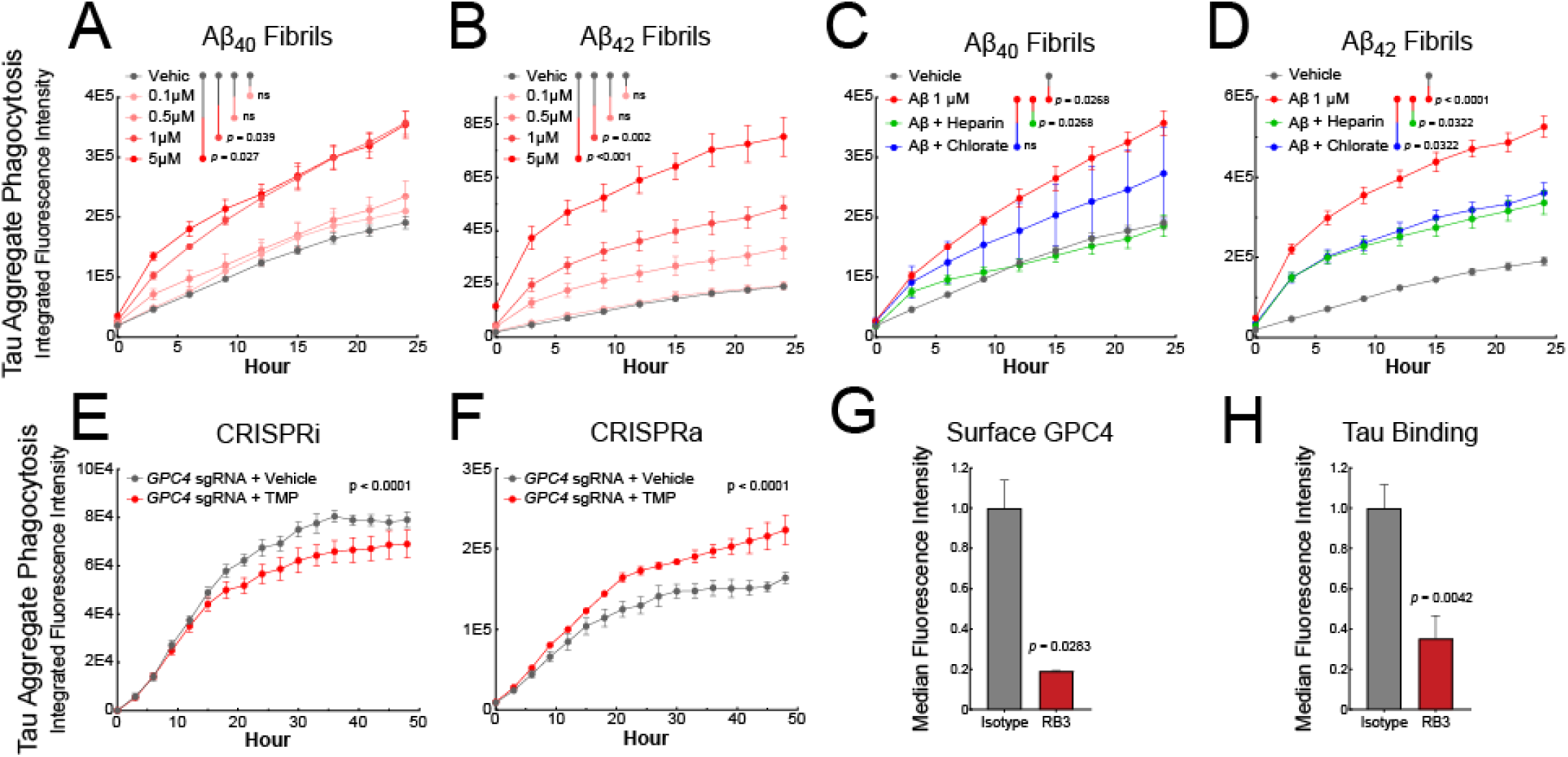
Heparan sulfate and GPC4 mediate tau phagocytosis in iTF Microglia. Phagocytosis of pHrodo red-labeled tau fibrils after pretreatment with Aβ_40_ (A) or Aβ_42_ (B) fibrils by iTF-Microglia. Phagocytosis of pHrodo red-labeled tau fibrils in the presence of heparan sulfate proteoglycan inhibitors heparin (100 µg/mL) or chlorate (25 mM) after pretreatment with Aβ_40_ (C) or Aβ_42_ (D) fibrils. The statistical analysis for experiments A – D were performed with a one-way ANOVA and Holm-Sidak multiple comparisons test. N = 4. Phagocytosis of pHrodo red-labeled tau fibrils using inducible CRISPRi (E) or inducible CRISPRa (F) iTF Microglia transduced with GPC4 sgRNAs. The CRISPRa and CRISPRi elements are activated by trimethoprim (50 nM). The statistical analyses for experiments E, F were performed with a paired t-test. N = 4. (G) Flow cytometry analysis of iTF Microglia measures the abundance of cell-surface GPC4 after treatment with α-GPC4 sdAb-Fc or isotype control antibodies for 24 hours. (H) Flow cytometry analysis of AF647-labeled tau fibrils binding to the iTF Microglia cell surface after pre-treatment with an α-GPC4 antibody. The cell-surface binding experiment was performed at 4°C. The statistical analyses were performed with a Student t-test. The data represent the means ± SEM.

We next tested whether a GPC4 function-blocking antibody can inhibit tau fibril phagocytosis. RB3 is an anti-GPC4 single domain antibody (sdAb) that was previously demonstrated to inhibit GPC4 activity resulting in increased potential of pluripotent stem cells to differentiate into midbrain dopaminergic neurons.^43^ We grafted RB3 onto a human IgG1 Fc scaffold to create a bivalent anti-GPC4 sdAb-Fc (**Figure S6B**). Treatment of iTF Microglia with 100 nM of RB3 for 24 h resulted in depletion of cell-surface GPC4 by ∼80% as measured by flow cytometry with a commercial polyclonal antibody, presumably via the increased intracellular trafficking and turnover of RB3-bound GPC4 (**Figure 5G**). This antibody treatment also led to a 65% reduction of tau-647 fibril binding to the iTF Microglia cell-surface in a cell-based binding experiment performed at 4°C (**Figure 5H**). Taken together, these data demonstrate that GPC4 mediates tau fibril binding and phagocytosis in iTF Microglia.

### Aβ fibrils lead to GPC4 shedding which promotes tau phagocytosis

GPC4 is a GPI-anchored HSPG that can exist in multiple proteoforms. To determine which GPC4 proteoform is most bioactive in mediating tau phagocytosis, we created three GPC4 genetic variants that encode 1) the natural cell-surface bound GPC4 (GPC4 WT), 2) a GPC4 variant lacking heparan sulfate (HS) chains due to ablated HS attachment sites (GPC4-ΔHS), and 3) a constitutively secreted GPC4 variant that lacks the GPI-anchor (GPC4-sec). We expressed these three variants in HEK cells and mouse microglia BV2 cells and measured their ability to internalize pHrodo-labeled tau fibrils. Overexpression of GPC4 WT enhanced tau fibril internalization by 76% in HEK293T cells and 25% in BV2 cells compared to NanoLuc (NLuc) expression control (**Figure 6A and S7A**). These data are consistent with the GPC4 CRISPRa experiment in iTF Microglia (**Figure 5F**). In contrast, GPC4-ΔHS failed to augment tau fibril phagocytosis suggesting a critical role for heparan sulfate chains in tau uptake as we previously reported.^35^ Surprisingly, we found that the constitutively secreted variant of GPC4, GPC4-sec, was an equally bioactive proteoform, increasing tau fibril internalization by 63% in HEK293T cells and 36% in BV2 cells. (**Figure 6A and S7A**). This suggests that both the membrane bound and soluble forms of GPC4 are bioactive in mediating tau internalization into cells, and require their heparan sulfate chains.

**Figure 6:**
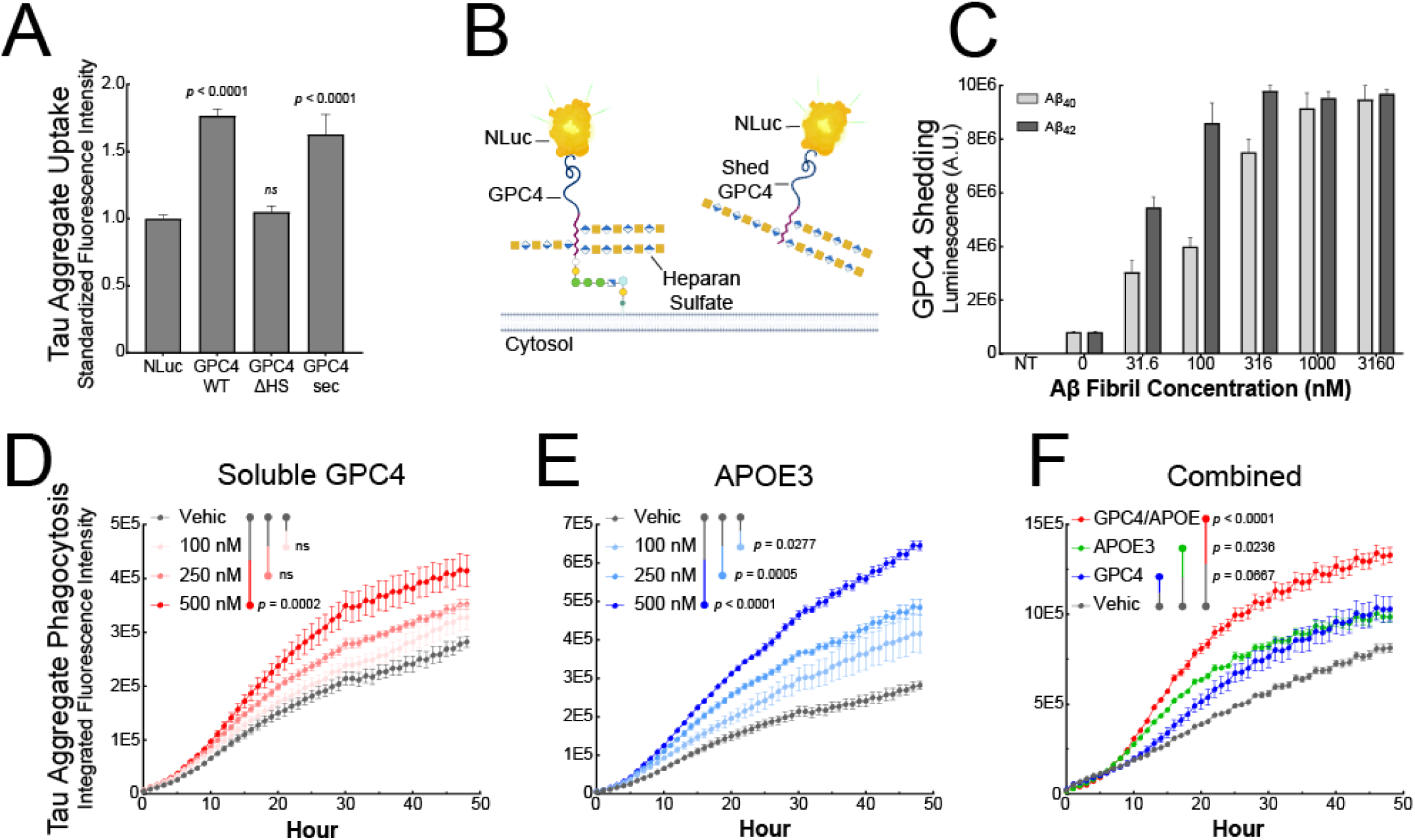
β-Amyloid fibrils lead to GPC4 shedding and APOE secretion which promote tau phagocytosis. (A) GPC4 WT, GPC4-ΔHS, GPC4-sec, or NLuc control plasmids were transiently transfected into HEK293T cells and tau aggregate-AF647 uptake was measured via flow cytometry. (B) Schematic model depicting NLuc-GPC4 fusion protein for tracking GPC4 shedding into the conditioned media via luminescence. (C) NLuc-GPC4 luminescence was measured in iTF Microglia conditioned media after a 24 h treatment with Aβ_40_ and Aβ_42_ fibrils. pHrodo-red labeled tau aggregates (50 nM) were preincubated with (D) soluble recombinant GPC4 or (E) soluble recombinant APOE3 and uptake was measured every hour for 48 h with an Incucyte SX5. (F) pHrodo-red labeled tau aggregates (50 nM) were preincubated with soluble recombinant GPC4 alone, APOE3 alone, or GPC4 + APOE3 (250 nM) and uptake was measured every hour for 48 h. The statistical analyses were performed with a one-way ANOVA and Holm-Sidak multiple comparisons test. N = 4. The data represent the means ± SEM.

During development, GPC4 is proteolytically shed from the cell-surface of astrocytes, and once released into the extracellular space, is known to promote the maturation of excitatory synapses.^44,45^ We next asked whether Alzheimer’s disease pathology can lead to GPC4 shedding from microglia thereby releasing soluble GPC4 into the extracellular space. To measure GPC4 shedding in iTF microglia, we constructed a luminescence reporter system by genetically fusing NLuc after the signal peptide of GPC4 (NLuc-GPC4) as has been previously reported.^44^ Luminescence can then be measured in the iTF Microglia conditioned media to quantify the shedding of GPC4 in various disease-relevant conditions (**Figure 6B**). We treated Nluc-GPC4 expressing iTF microglia with Aβ_40_ or Aβ_42_ fibrils for 24 h and observed a dose-dependent increase, up to 10-fold, in shed GPC4 as measured by luminescence in the conditioned media (**Figure 6C**). This effect was not observed with scrambled Aβ nor when the experiment was performed in the presence of sodium azide, a metabolic inhibitor of ATP production, suggesting that this phenomenon is amyloid-specific and requires cell metabolism (**Figure S7B and S7C**).

Given the possibility that Aβ aggregates can result in the shedding of GPC4, we next asked if soluble GPC4 is sufficient to promote tau phagocytosis into iTF Microglia. We co-applied pHrodo-labeled tau fibrils with increasing doses of recombinant, soluble GPC4 lacking a GPI-anchor. We observed that soluble GPC4 was sufficient to dose-dependently augment tau phagocytosis (**Figure 6D**), consistent with the hypothesis that extracellular GPC4 may potentiate tau aggregate internalization.

APOE is known to potentiate tau pathology and increase tau aggregate internalization into neurons,^22,46,47^ as well as interact with astrocyte-secreted GPC4 to drive tau hyperphosphorylation.^17^ Since APOE was significantly enriched in our surfaceomics dataset for microglia treated with Aβ fibrils (**Figure 1A and 1B**), we next investigated whether APOE can alter microglial phagocytosis of tau aggregates. We confirmed that microglia secrete APOE after stimulation with Aβ_40_ or Aβ_42_ fibrils using an APOE ELISA on conditioned media (**Figure S7D**). We then asked whether soluble APOE is sufficient to potentiate tau aggregate phagocytosis in microglia. We co-applied mammalian-derived, lipidated APOE3^48^, the most common allele in human populations, with tau-pHrodo fibrils. We observed a significant and dose-dependent increase in tau fibril phagocytosis with the addition of APOE3 (**Figure 6E**). Lastly, we asked whether the combined addition of GPC4 and APOE3 could further potentiate tau phagocytosis. We applied tau-pHrodo fibrils with either GPC4 alone, APOE3 alone, or both GPC4 and APOE3. We observed that the addition of GPC4 and APOE3 together significantly potentiated tau aggregate phagocytosis relative to treatment with GPC4 or APOE3 alone (**Figure 6F**). The addition of N-terminal APOE3 lacking the lipid-binding C-terminal domain, did not result in any augmentation of tau phagocytosis (**Figure S7E**). Taken together, these data suggest that GPC4 and APOE3 work in concert to drive tau aggregate phagocytosis in microglia.

### Tau binds to GPC4 on the cell-surface

We next investigated whether GPC4 mediates tau binding to the cell-surface. We expressed either GPC4 WT or heparan sulfate deficient GPC4-ΔHS in HEK cells and measured their ability to facilitate cell-surface binding of tau-647 fibrils at 4 °C. GPC4 WT expression resulted in ∼3.5-fold more tau fibril binding to the cell-surface relative to the NLuc expression control vector (**Figure S8A**) confirming that GPC4 is sufficient to mediate cell-surface binding of tau fibrils. In contrast, GPC4-ΔHS minimally augmented tau cell-surface binding, further corroborating the importance of the heparan sulfate chains in facilitating interactions with tau. Additionally, when HEK293T cells were transfected with a secreted form of NLuc-GPC4 WT, we successfully pulled down NLuc-GPC4 from the conditioned media using monomeric tau-biotin immobilized on streptavidin beads, as measured by luminescence (**Figure S8B**). This interaction was significantly enhanced by the addition of APOE3, suggesting that tau, GPC4, and APOE3 may form a multiprotein complex.

However, we were unable to detect direct biophysical interactions between tau, GPC4, and APOE3 when using recombinant purified proteins. Biolayer interferometry (**Figure S8C**), protein pull-down assays (**Figure S8D and S8E**), and lysine crosslinking with disuccinimidyl sulfoxide (DSSO) (**Figure S8F**) did not reveal direct protein-protein interactions. These data indicate that the cellular expression of GPC4 WT is sufficient to mediate cell-surface binding of tau, but that purified recombinant GPC4 is insufficient to bind tau in solution. This suggests the involvement of an additional, yet unidentified, co-factor or cellular process in mediating interactions between tau, GPC4 and APOE3.

### Microglia-derived GPC4 and APOE interact to amplify tau seeding in iPSC Neurons

According to the prion hypothesis of Alzheimer’s disease, misfolded tau can propagate from one neuron to another, spreading tau pathology across the brain. The process involves the release of misfolded tau from a donor cell, uptake by a recipient cell, and subsequent templating where the misfolded tau induces normal tau protein to misfold and aggregate. Our data demonstrate that microglia release GPC4 and APOE in response to Aβ fibrils which, in turn, potentiates tau internalization. Given that soluble factors can act *in trans* and not only *in cis*, we asked whether soluble GPC4 and APOE can augment tau fibril uptake and pathological seeding in neurons.

We co-applied tau-647 fibrils to SH-SY5Y cells, a human neuroblastoma cell line, along with GPC4 and APOE3, either alone or in combination, and measured internalization via flow cytometry 16 hours later. We found that GPC4 and APOE3 alone increased tau aggregate internalization by 1.5-fold and 1.6-fold, respectively, while their co-application increased tau aggregate uptake by 2.35-fold (**Figure 7A**). These findings are consistent with our previous results showing that GPC4 and APOE3 together enhance tau aggregate phagocytosis in iTF Microglia (**Figure 6F**). To evaluate whether tau fibrils, GPC4, and APOE3 colocalize within treated cells, we performed confocal microscopy on living SH-SY5Y cells. Tau-647 fibrils were co-treated with GPC4-546 and APOE3-488 for 16 h, followed by trypsinization to remove cell-surface bound protein, replating, and confocal imaging. We observed colocalization of tau with both GPC4 and APOE3 (**Figure 7B**), suggesting that these proteins are localized within the same cellular compartments after internalization.

**Figure 7:**
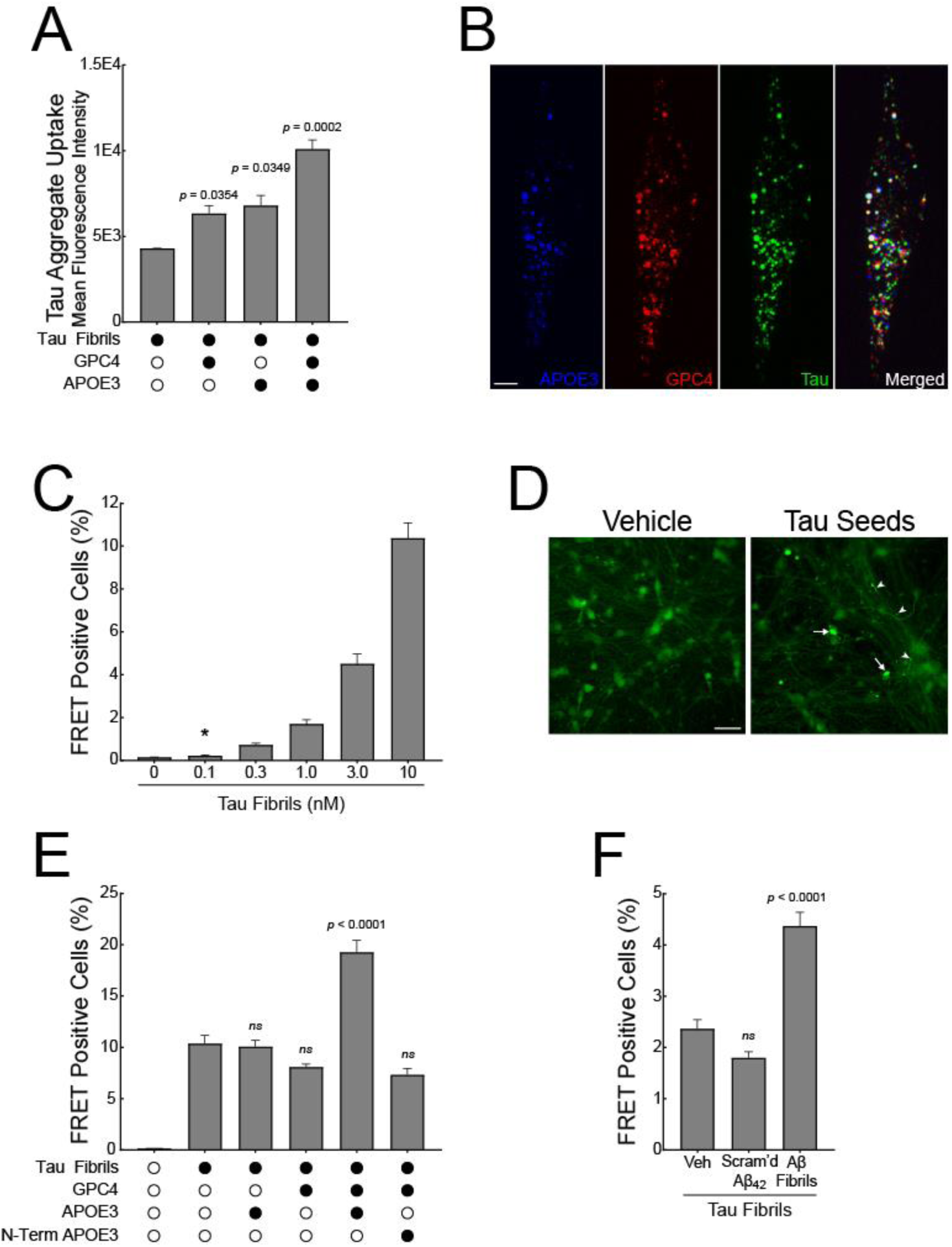
GPC4 and APOE3 amplify tau seeding in iNeuron FRET biosensors. (A) Uptake of tau-pHrodo red fibrils (50 nM) with or without GPC4 or APOE3 (250 nM) measured at 16 h in SK-N-AS cells. (B) SK-N-AS cells were co-incubated with tau-647 fibrils, GPC4-546 or APOE3-488 for 16 hours, trypsinized, replated, and then imaged on a confocal microscope 4 h later. Scale bar = 10 µM. (C) iNeuron tau FRET biosensors were treated with varying doses of full-length tau fibrils for 7 days prior to measuring intraneuronal tau pathology by FRET flow cytometry. The assay is linear and statistical significance is first reached at a tau fibril concentration of 0.1 nM. (D) Confocal images of iNeuron tau FRET biosensors treated with vehicle or 10 nM tau fibrils for 7 days. The arrows mark tau aggregates within the neuron soma and the arrowheads mark tau aggregates within neuron processes. Scale bar = 40 µM. (E) iNeuron tau FRET biosensors were treated with tau fibrils (10 nM) with or without GPC4, APOE3, or N-terminal APOE3 (50 nM) for seven days prior to FRET flow cytometry. (F) iNeuron tau FRET biosensors were treated with Aβ_40/42_-primed microglia conditioned media containing 10 nM tau fibrils for 7 days prior to FRET flow cytometry. The statistical analyses were performed with a one-way ANOVA and Holm-Sidak multiple comparisons test. N = 4. The data represent the means ± SEM.

Next, we asked whether the co-application of GPC4 and APOE, along with tau fibrils, could potentiate tau aggregation in a neuronal culture system. Previously, we generated a HEK293T tau FRET biosensor cell line that specifically detects tau seeds in the femtomolar range.^49,50^ Here, we report the development of a more physiologically relevant biosensor based on iPSC neurons stably expressing the tau repeat domain (RD) with the disease-associated P301S mutation fused to either mClover or mApple. At baseline, the biosensor reporter proteins are monomeric. However, the application of exogenous tau seeds induces tau reporter protein aggregation, resulting in a FRET signal that can be measured with microscopy or flow cytometry. To validate this system, we applied a dose range of unlabeled, recombinant 2N4R tau fibrils to iNeuron biosensor cells without the addition of liposomes. After 7 days, the tau seeds induced a dose-dependent increase in intraneuronal tau pathology, with 100 pM of exogenous tau seeds initiating tau aggregation, demonstrating that the iNeuron biosensor system is sensitive and quantitative (**Figure 7C and 7D**).

We then applied tau seeds (10 nM) along with recombinant GPC4 and APOE3 (50 nM) and measured tau pathology after 7 days. Tau seeds alone resulted in FRET positivity in approximately 10% of the iNeuron biosensors. This response was not augmented by the addition of either GPC4 or APOE3 alone. However, the co-application of tau seeds with both GPC4 and APOE3 resulted in a doubling of FRET positive neurons (**Figure 7E**). These data suggest that GPC4 and APOE3 work together to augment tau seeding in iNeurons.

Finally, we induced the secretion of GPC4 and APOE3 by priming iTF microglia with 1 µM Aβ_40/42_ fibrils for 24 h and then harvested the conditioned media. We mixed the conditioned media with 10 nM tau fibrils and applied the mixture to the iNeuron biosensors for 7 days. Aβ-primed microglia conditioned media significantly potentiated tau seeding compared to vehicle-primed or scrambled Aβ_42_-primed cells (**Figure 7F**), confirming that Aβ fibrils induce microglial secretion of factors that enhance tau pathology in neurons.

## DISCUSSION

Aβ and tau are critical proteins in the pathogenesis of AD, but the mechanisms by which they interact to promote disease remain unclear. Our study delineates a role for microglia in mediating the interaction between Aβ and tau pathology through the expression of the GPI-anchored heparan sulfate proteoglycan, GPC4. In concert with APOE3, GPC4 enhances the entry of tau seeds into both microglia and neurons, with neuronal entry facilitating subsequent tau pathology.

To catalogue the human iPSC microglia surfaceome in response to Aβ, we first employed a rapid and sensitive cell-surface proteomic strategy using WGA-HRP.^27^ Our proteomics data revealed that Aβ-exposed microglia upregulate heparan sulfate proteoglycans and the heparan-sulfate binding protein, APOE. Using flow cytometry, we validated that microglia selectively upregulate heparan sulfate and GPC4 when treated with amyloids but not cognate monomer or innate immune activators. Additionally, iPSC astrocytes, iPSC neurons, and immortalized cell lines, minimally altered their cell-surface heparan sulfate when exposed to Aβ fibrils. Thus, the process of Aβ-induced microglia priming may exhibit both substrate and cell-type specificity.

We validated these findings in human AD post-mortem tissue. Specifically, we found that AD microglia express GPC4 in proximity to Aβ plaques and that overall microglia GPC4 expression correlates with Aβ plaque burden, but not neurofibrillary tangle burden. These findings are reminiscent of a recent report demonstrating that the lipid-processing enzyme *ACSL1* is enriched in microglia near Aβ plaques in AD brain tissue.^51^ Our study does not address the possibility that other CNS cell types may express GPC4. However, another recent effort found that astrocyte-secreted GPC4 preferentially interacts with APOE4 over APOE3 and APOE2 to promote tau hyperphosphorylation.^52^ Notably, their data demonstrate that AD patients harboring two copies of APOE4 have a modest elevation of GPC4 (∼0.25-fold) while AD patients homozygous for APOE3 or APOE2 have similar astrocyte GPC4 expression to age-matched controls. Therefore, the astrocyte GPC4 upregulation appears to be due to APOE4 alleles and not Aβ pathology. In our study, we observed that AD-associated microglia upregulate GPC4 2.2-fold in an Aβ-plaque dependent manner. Additionally, three out of four AD subjects were APOE3 homozygous, and the fourth subject was APOE3/4 heterozygous (Table 1), implying that microglia GPC4 expression is independent of the APOE4 allele. Thus, consistent with our cell culture results, it appears that microglia may be the dominant cell type responding to Aβ by upregulating GPC4. It is notable that neither GPC4 nor any member of the glypican family have been previously associated with microglia biology. We predict that a complex interplay between glial GPC4 expression, APOE genotype, and amyloid pathology in AD may exist that warrants further examination.

To investigate whether GPC4 modulates Aβ-associated deficits *in vivo*, we used a *Drosophila* amyloidosis model that expresses human Aβ_42_ pan-neuronally.^41^ This model displays decreased climbing activity and early lethality. We found that glial-specific over-expression of *Dlp*, the *Drosophila* ortholog of GPC4, exacerbated Aβ-mediated toxicity, whereas the knockdown of glial *Dlp* rescued Aβ-mediated toxicity. Importantly, these phenotypes were not recapitulated when *Dlp* expression was modulated in a non-transgenic *Drosophila* line.

These findings demonstrate that glial GPC4 interacts with Aβ to enhance toxicity *in vivo*, but the mechanisms mediating these phenotypes remain unclear.

Our prior work establishes that cell-surface HSPGs mediate the internalization of proteopathic amyloids including tau and synuclein, facilitating the prion-like spread of pathology from cell to cell.^35–37,53,54^ We therefore reasoned that GPC4 may similarly mediate the uptake of tau aggregates. Indeed, other groups have demonstrated that GPC4 enables the internalization of Aβ in neural stem cells^55^, tau oligomers in astrocytes^56^, or tau monomer in neurons.^52^ Here, we found that Aβ primes microglia to express cell-surface heparan sulfate and GPC4. Using a combination of pharmacologic-, genetic-, and antibody-based inhibition, we found that GPC4 regulates the binding and phagocytosis of tau aggregates into microglia, consistent with the established role of HSPGs as a receptor for tau aggregates. Unexpectedly, however, we also found that Aβ leads to GPC4 cell-surface shedding, resulting in soluble, extracellular GPC4 proteoforms. Soluble GPC4 retained bioactivity in facilitating tau phagocytosis suggesting that the cell-surface anchored proteoform of GPC4 may not be the only pathogenic form. We do not yet know the mechanisms of Aβ-induced GPC4 shedding or the mechanisms by which soluble GPC4 mediates tau aggregate internalization.

Classically, GPC4 is an established effector of CNS development. For example, astrocyte-derived GPC4 and GPC6 induce the formation of functional synapses in developing mouse retinal ganglion neurons.^45^ In rats, GPC4 is localized to neuronal pre-synaptic membranes where it regulates the development of excitatory neurons.^57^ In the developing mouse cerebral cortex, GPC4 is expressed in neural precursor cells to regulate FGF2 signaling and cortical neurogenesis.^58^ Here, we propose a role for GPC4 in microglia biology in the context of neurodegeneration. As GPI-anchored proteins, glypicans possess the unique ability to detach and reinsert themselves into the plasma membrane, enabling lateral cell-to-cell translocation and ligand gradient formation, a process referred to as *hopping*.^59^ During development, hopping is exploited to generate morphogen gradients for the proper patterning of CNS topology. For example, in *Drosophila*, the extracellular distribution of secreted Wnts is established via the GPC4 ortholog, *Dlp*, which facilitates short- and long-range signaling gradients to ensure proper formation of the germarium and wing discs. Further, *Dlp* shields Wnt’s lipid moiety from the aqueous environment effectively solubilizing the ligand and permitting its spread to more distant sites.^60^ It is intriguing to speculate that, in neurodegeneration, GPC4 hopping may be hijacked to traffic pathologic proteins, such as tau or inflammatory cytokines, from amyloid plaques to distant sites via diffusion-based gradients as has been observed for synthetic GPI-anchored ligands.^59^ Future studies should test whether this mechanism can explain the network diffusion models of disease spread in neurodegeneration.^7,61,62^

Coincident with this hypothesis, we observed that the lipid-bearing protein, APOE is also secreted by Aβ-treated microglia. APOE is a heparan sulfate binding protein^63^, and APOE3 and APOE4 facilitate tau aggregate entry into neurons, whereas the protective Christchurch mutation abrogates tau uptake.^47,64^ Astrocytes are the primary cell type to express APOE in the healthy human brain. However, in AD, astrocytes downregulate and microglia upregulate APOE production.^65–67^ We therefore hypothesized that microglia-secreted GPC4 and APOE may work together to facilitate tau internalization and pathological seeding. We found that APOE3 is sufficient to stimulate tau phagocytosis in microglia, and the combination of GPC4 plus APOE3 resulted in synergistic tau uptake. Although phagocytosis of tau aggregates in microglia could be a neuroprotective function that eliminates pathologic tau aggregates, numerous studies now suggest that microglia tau phagocytosis can instead accelerate the spread of tau pathology via extracellular vesicles.^19,68–70^ We have yet to explore the impact of GPC4 on the formation of extracellular vesicles.

To investigate if soluble GPC4 and APOE together enhance tau seeding in neurons, we developed a novel iPSC neuron tau FRET biosensor line that revealed that the co-addition of GPC4 and APOE3 enhanced pathological tau seeding. This effect was absent when GPC4 or APOE3 were applied individually, suggesting that there is a synergistic relationship between GPC4 and APOE3 for tau uptake and seeding. Furthermore, conditioned media from Aβ-treated microglia potentiates tau seeding, establishing a role for microglia secreted factors as contributors to tau pathology. We therefore propose that microglia are particularly well-suited to respond to Aβ via a toxic GPC4 / APOE secretion axis and that this microglial axis may accelerate neurodegeneration by amplifying tau pathology and spread in neurons. These observations are important for the development of therapeutic agents that target the intersection of Aβ and tau pathology.

### Limitations of the Study

Our data demonstrate a biological interaction between Aβ, GPC4, APOE3, and tau. However, we have yet to define the mechanisms by which Aβ induces GPC4 expression. Additionally, we have not yet determined whether other APOE alleles impact their interaction with GPC4. Given that APOE allele variants alter APOE’s affinity for heparan sulfate^71,72^, we predict that these variants may modulate GPC4’s impact on tau pathology. Further, we used a *Drosophila* model to investigate the role of glial GPC4, but our genetic manipulations do not differentiate between astrocytes and cortex glia (the fly analog of microglia), and thus we are unable to make claims about microglia-specific versus astrocyte-specific effects in these in vivo studies. Finally, *Drosophila* tau is not known to spontaneously aggregate, and therefore it is unlikely that the effects of GPC4 on Aβ-mediated toxicity operate via tau pathology. This implies that there may be other, non-tau, mediated effects governing GPC4-amyloid toxicity.

## METHODS

### Human iPSC-derived iTF-Microglia cell culture and differentiation

We generated iPSC-derived iTF Microglia using a protocol established by Dräger et al., 2022^32^. Human iPSCs (WTC11, Coriell Cat. No. GM25256) with stably integrated doxycycline-inducible transcription factors (CEBPa, CEBPb, IRF5, IRF8, MAFB, PU1) were maintained according to established protocols^32^. For differentiation, day 0 iPSCs were seeded onto plates coated with Matrigel and poly-d-lysine in Essential 8 Medium (Gibco; A1517001) supplemented with 10 µM ROCK inhibitor and 2 µg/mL doxycycline (Clontech; 631311). On day 2, the media was changed to Advanced DMEM/F12 Medium (Gibco; 12634-010) supplemented with 1 × Antibiotic-Antimycotic (Anti-Anti) (Gibco; 15240-062), 1 × GlutaMAX (Gibco; 35050-061), 2 μg/mL doxycycline, 100 ng/mL Human IL-34 (Peprotech; 200-34) and 10 ng/mL Human GM-CSF (Peprotech; 300-03). On day 4, 6, and 8 the media was replaced with Advanced DMEM/F12 supplemented with 1 × Anti-Anti, 1 × GlutaMAX, 2 μg/mL doxycycline, 100 ng/mL Human IL-34, 10 ng/mL, Human GM-CSF, 50 ng/mL Human M-CSF (Peprotech; 300-25), and 50 ng/mL Human TGF-β1 (Peprotech; 100-21C). On day 8, the differentiated iTF-Microglia were used for experimentation. For CRISPRi and CRISPRa experiments, iTF iPSCs harboring trimethoprim-inducible CRISPRi or CRISPRa machinery^32^ were used and maintained in 50 nM trimethoprim (MP Biomedical; 195527) starting on day 0 and continuing through the remainder of differentiation.

### Human iPSC Neuron Cell culture, Differentiation, and Generation of FRET Biosensors

Pre-differentiation and differentiation of excitatory glutamatergic neurons was performed as previously described.^73^ Briefly, iPSCs were split into matrigel coated plates in pre-differentiation medium (KnockOut DMEM/F12 with 1× MEM non-essential amino acids, 1× N2 Supplement [Gibco/Thermo Fisher Scientific, 17502-048], 10 ng/mL of BDNF [PeproTech, 450-02], 10 ng/mL of NT-3 [PeproTech, 450-03], 1 μg/mL of mouse laminin [Thermo Fisher Scientific, 23017-015], and 2 μg/mL of doxycycline [Takara; 631311]). 10 nM ROCK inhibitor was added on the initial passage day (Day -3) and omitted thereafter. After a total of three days in pre-differentiation media (day 0), cells were re-plated on poly-D-lysine coated BioCoat plates (Corning) in BrainPhys media containing 0.5× N2 supplement, 0.5× B27-VA supplement, 10 ng/mL NT-3, 10 ng/mL BDNF, 1 µg/mL mouse laminin, and 2 µg/mL doxycycline. On day 3, media was replaced with complete BrainPhys media without doxycycline. Subsequently, half media changes were performed weekly unless stated otherwise. For uptake experiment, iNeurons were treated on day 7. For seeding experiments, iNeurons were treated on day 3 and harvested for flow cytometry on day 10.

To produce tau FRET biosensor iPSCs, lentivirus encoding tauRD (244 – 368)-mClover and mApple harboring the P301S mutation was transduced into the previously established CRISPRi-i3N human iPSC line^73^ (Coriell GM29371). A polyclonal population of dual-positive TauRD(P301S)-Clover/Apple biosensors were sorted using a BD FACSAria Fusion and expanded for further use.

### Human iPSC Astrocytes

Differentiation of iAstrocytes was performed as previously described.^74^

### Cell Culture, Transfection, and Viral Infection of Cells

HEK293 cells (ATCC, CRL-1573) were cultured in high-glucose DMEM with 10% FBS and penicillin (100 U/ml)-streptomycin (100 µg/ml). SH-SY5Y (ATCC, CRL-2266) cells were cultured in advanced DMEM/F12 (Gibco) supplemented with 10% FBS, penicillin (100 U/ml), and streptomycin (100 µg/ml). Cells were incubated in a humidified atmosphere of 5% CO_2_ at 37 °C. HEK293 cells were transfected with plasmids using Lipofectamine 3000 (Thermo Fisher Scientific; L3000008) or transduced with lentivirus encoding GPC4 WT, GPC4-secreted, GPC4-ΔHS, GPC4-secreted-ΔHS, NLuc-GPC4 WT, or NLuc-GPC4-secreted. All GPC4 DNA sequences were synthesized by Twist Biosciences and cloned into FM5 vector using a Gibson assembly method.

### Protein Acquisition, Expression, Purification, Fibrillization, and Labeling

Recombinant Aβ_40_ (A-1001-2), Aβ_42_ (A-1002-2), scrambled Aβ_42_ (A-1004-1), and α-synuclein (S-1001-1) were purchased from rPeptide. Aβ_40_ and Aβ_42_ peptides were dissolved at 5 mg/mL in hexafluoroisopropanol and evaporated into a peptide film. The peptide films were resuspended at 10 mg/mL in DMSO, vortexed, and sonicated for 5 min, then diluted to 0.2 mg/mL in 10 mM sodium phosphate buffer, pH 7.4. The solution was shaken at 900 rpm at 37 °C for 72 h (Aβ_40_) or 120 h (Aβ_42_) to form fibrils. Scrambled Aβ_42_ (A-1004-1) was reconstituted at 0.2mg/mL in PBS and stored at -80 C° until further use. α-Synuclein was fibrillized by dissolving the lyophilized protein at 2mg/mL in PBS at 900 rpm at 37 °C for 72 h; α-Synuclein monomer was maintained by immediately storing the freshly reconstituted protein at -80 °C without shaking. Full length 2N4R tau with a C-terminal polyhistidine tag in the pet28b plasmid was expressed and prepared as previously described^75^ from BL21-Gold (DE3) competent cells. Tau was purified on a Ni-NTA column and eluted in 1× PBS. To induce fibrillization of tau monomer, 8 μM tau was incubated at 37 °C in 10 mM HEPES, 100 mM NaCl, and 8 μM heparin for 72 h without agitation. Tau and magnetic beads (Spherotech; AM-10-10) were conjugated to pHrodo-red dye or AlexaFluor 647 (ThermoFisher; P36600 and A20006) by buffer exchanging the substrates into 100 mM sodium bicarbonate buffer, adding pHrodo red dye at a final molar ratio of 5:1 (dye : substrate), and incubating at RT for 15 min. Excess dye was quenched by adding glycine at a final concentration of 100 mM for 15 min and then buffer exchanging via dialysis into PBS. For protein pull-down assays, full-length tau-biotin was purchased from rPeptide (T-1114-1). Mammalian-derived and heparan-sulfated GPC4-WT was purchased from Acro (GP4-H52H3).

For APOE expression and purification, human FL APOE3 and N-terminal APOE3 (1 – 216) DNA sequences were generated by Twist Bioscience and cloned into pcDNA3.4 using a Gibson assembly method. Transient transfection of plasmid DNA was done in Expi293F cells at 37 °C. Cells were harvested on day 7 by centrifugation at 4000 x g for 20 min and the supernatant was filtered through a 0.45 µM sterile filter. The supernatant was incubated with 1mL (per 30 mL of supernatant) Ni-NTA resin (Cytiva; 17371203) and imidazole to a final concentration of 5 mM for 1 h at 4 °C with end-over-end mixing. The resin-media mixture was loaded on a chromatography column and washed with 5 resin volumes of wash buffer (PBS pH 8.0, 5 M NaCl, 50 mM K_3_PO_4_, 10 mM imidazole) for a total of two times. APOE3 was then eluted in 1 resin volume of elution buffer (0.5 M NaCl, 50 mM K_3_PO_4_, 1 M imidazole) in a total of 3 elution fractions. The purity of each fraction was assessed by SDS-PAGE and fractions > 90% pure were pooled and buffer exchanged into PBS using a 10 kDa MWCO centrifugal filter (Amicon; UFC9010). The protein concentration was determined by bicinchoninic acid (BCA), and the protein was diluted to a final concentration of 8 µM and stored at -80 °C.

### iTF Microglia Cell-Surface Labeling

For cell-surface proteomic studies, the iTF Microglia cell-surface was labeled with cell-tethered WGA-HRP according to the protocol established in Kirkemo et al., 2022^27^. In brief, cells were lifted using Versene (Gibco; 15040066) for 10 min at 37 °C. Cells were diluted in DPBS and pelleted by centrifugation at 1500 g for 5 minutes and resuspended in DPBS pH 6.5. Cell surfaces were labeled by incubating cells with 0.5 µM WGA-HRP for 5 minutes on ice, then adding 500 µM biotin tyramide, and finally adding 1 mM of H_2_O_2_. The mixture was incubated at 37 °C for 2 minutes. The reaction was quenched with 10 mM Sodium Ascorbate/1 mM Sodium Pyruvate followed by two additional washes in the same buffer prior to a final wash in DPBS. Cells were pelleted and flash frozen before further processing.

### Proteomic Preparation for Surface-Labeled iTF Cells

Frozen cell pellets were thawed and processed for LC-MS/MS using a preOmics iST kit (P.O. 00027). Briefly, cell pellets were lysed in 2x RIPA buffer (Millipore Sigma; 10-188) containing protease inhibitors (cOmplete, Mini, EDTA-free Protease Inhibitor Cocktail; Millipore Sigma; 11836170001) and 1.25 mM EDTA. Cells were further disrupted via sonication. Biotinylated proteins were pulled down with NeutrAvidin-coated agarose beads (ThermoScientific; 29204) for 1 hour at 4 °C. Beads were transferred to Poly-Prep chromatography columns (Bio-Rad) and sequentially washed with 1x RIPA buffer, high-salt PBS (PBS pH 7.4, 2 M NaCl), and denaturing urea buffer (50 mM ammonium bicarbonate, 2 M Urea). From the PreOmics iST kit, 50 µL of the provided LYSE solution was added to the slurry and the mixture was incubated at 55 °C for 10 min with shaking. The provided enzyme mixture (Trypsin and LysC) was resuspended in 210 µL of RESUSPEND buffer, mixed, and 50 µL was added to the slurry. Samples were allowed to mix at 500 rpm for 90 minutes at 37 °C, before being quenched with 100 µL of STOP solution. Samples were desalted using the provided Preomics columns and wash buffers per the manufacturer’s instructions. Peptides were eluted with 2Χ 100 µL of ELUTE, dried, and resuspended with 2% acetonitrile and 0.1% TFA. Peptides were quantified using Pierce Quantitative Colorimetric Peptide Assay (Thermo Fisher Scientific, 23275).

### LC-MS/MS and Data Analysis

Liquid chromatography and mass spectrometry were performed as described previously^27^. Briefly, 200 ng of samples were separated over a 90-minute linear gradient of 3-35% solvent B (Solvent A: 2% acetonitrile, 0.1% formic acid, solvent B: acetonitrile, 0.1% formic acid) on either an Aurora Ultimate CSI 25cm×75µm C18 (Ionopticks) or a PepSep XTREME 25 cm x 150 µm (Bruker) UHPLC column using a nanoElute UHPLC system (Bruker), and injected into a timsTOF pro mass spectrometer (Bruker). Data-dependent acquisition was performed with parallel accumulation-serial fragmentation (PASEF) and trapped ion mobility spectrometry (TIMS) enabled with 10 PASEF scans per topN acquisition cycle. For database searching, peptides were searched using MaxQuant’s (Version 2.6.1) Andromeda search engine against the plasma membrane annotated human proteome (Swiss-prot GOCC database, June 3, 2020 release). Enzyme specificity was set to trypsin + LysC with up to two missed cleavages. Cysteine carbamidomethylation was set as the only fixed modification; acetylation (N-term) and methionine oxidation were set as variable modifications. The precursor mass error tolerance was set to 20 PPM and the fragment mass error tolerance was set to 0.05 Da. Data was filtered at 1% for both protein and peptide FDR and triaged by removing proteins with fewer than 2 unique peptides. GPC6 was identified via a single peptide and was thus excluded from all proteomic analyses with the exception of Figure 1E where it was included to compare to other glypicans. All mass spectrometry database searching was based off of at least three biological replicates. Biological replicates underwent washing, labeling, and downstream LC-MS/MS preparation separately. Perseus was used to analyze label-free quantitation data generated in MaxQuant. All peak areas were log2(x) transformed and missing values were imputed separately for each sample using the standard settings (width of 0.3, downshift of 1.8). Significance was based off of a standard unpaired Student t test with unequal variances across all replicates. Reported peak area values represent the averages of all replicates. For representation of the data in figures, a Z-score was computed and is defined as (LFQ Area - Mean LFQ Area)/Standard Deviation. Protein IDs that were not annotated to be secreted or expressed extracellularly were removed. Heatmaps comparing expression levels between donors were generated with heatmapper.ca using an average linkage clustering method with Euclidean distance.

### Immunocytochemistry

All iTF-Microglia were grown on µ-slides (Ibidi; 80804) for imaging studies. For immunocytochemistry of microglia markers, day 8 iTF-Microglia were washed with PBS solution, fixed with 4% PFA, and blocked with 10% normal goat serum in PBS. Cells were stained with anti-IBA1 rabbit primary antibody (1:50; Cell Signaling Technology), anti-PU.1 rabbit primary antibody (1:100), and anti-TMEM119 mouse primary antibody (1:100) overnight at 4 °C followed by secondary labeling with Alexa Fluor 488 anti-mouse or Alexa Fluor 546 anti-rabbit (1:2000) and Hoechst 33342 (Invitrogen; H1399) for 1 h at RT. For the intracellular antigens IBA1 and PU.1, cells were permeabilized in 0.25% Triton X-100 prior to the addition of primary antibodies. For glypican staining, cells were treated with 1 µM Aβ_40_ or Aβ_42_ fibrils for 24 hours, washed three times in PBS, and fixed as described earlier. Cells were then blocked with 10% NGS and stained with GPC1 (1:100; Abcam), GPC2 (1:100; Invitrogen), GPC3 (1:200; Invitrogen), GPC4 (1:100; ProteinTech), GPC5 (1:200; R&D Systems), and GPC6 (1:100; Bioss) overnight at 4 C° followed by secondary labeling with Alexa Fluor 488 anti-mouse or Alexa Fluor 647 anti-rabbit (1:2000) and Hoechst 33342.

### Human Brain Tissue and Immunohistochemistry

Sixteen human brain tissue samples were obtained from the Neurodegenerative Disease Brain Bank at the University of California, San Francisco. Prior to autopsy, patients or their surrogates provided informed consent for brain donation, in keeping with the guidelines put forth in the Declaration of Helsinki. Neuropathological diagnoses were made following consensus diagnostic criteria^76^ using previously described histological and immunohistochemical methods.^77–79^ Cases were selected based on neuropathological diagnosis. Healthy control tissues were obtained from individuals without dementia who had minimal age-related neurodegenerative changes. Formalin-fixed blocks of the middle frontal gyrus were cut from coronal slabs and embedded together into tissue arrays in paraffin wax. Each tissue array contained four small tissue blocks with one tissue block from each histopathologic category (amyloid-/ tau-, amyloid+/tau-, amyloid-/tau+, and amyloid+/tau+). For immunofluorescence, tissue arrays were sectioned at 8 µm using a rotary microtome. The case details are listed in Table 1.

To reduce the autofluorescence of human brain tissue, glass-mounted sections were photobleached for 72 h using a multispectral LED array in a cold room^80^. The sections were baked at 65 °C for 30 minutes and deparaffinized and rehydrated followed by antigen retrieval in an autoclave at 120 °C for five minutes using 10 mM citrate buffer (pH 6). The tissue sections were permeabilized in PBS containing 0.25% Triton X-100 (PBS-T) and blocked in 10% normal goat serum for 1 hour. Primary antibodies were diluted in PBS-T and 10% normal goat serum and applied to the slides overnight at RT. The sections where then washed in PBS-T and secondary antibodies diluted 1:500, with or without DAPI, were applied at RT for two h. For experiments requiring amyloid plaque fluorescent staining, Amylo-Glow RTD (Biosensis; TR-300-AG) was applied for 10 minutes per the manufacturer’s directions after the secondary antibody application. The sections were washed with PBS and coverslipped with Fluoromount-G (ThermoFisher; 00-4958-02). For quantification of amyloid plaque and neurofibrillary tangle burden, tissue sections were deparaffinzed, peroxidase-blocked with 3% H_2_O_2_ in methanol for 30 minutes, and, for amyloid plaque staining, pre-treated with 88% formic acid for 6 minutes. Antigen retrieval and antibody staining was performed as described above. DAB reactions were developed by applying Vectastain ABC Elite (Vector Laboratory; PK-6200) followed by 0.05% diaminobenzidine and counterstaining with hematoxylin. Antibodies used in this study include: anti-IBA1 guinea pig primary antibody (1:250; Synaptic Systems), anti-Glypican 1 rabbit primary antibody (1:50; Abcam;), anti-Glypican 2 rabbit primary antibody (1:50; Invitrogen), anti-Glypican 4 rabbit primary antibody (1:50; ProteinTech), anti-Glypican 6 rabbit primary antibody (1:50; Bioss Antibodies), anti-Aβ mouse primary antibody (DE2; 1:500; Millipore Sigma), anti-phosphorylated tau (CP13; 1:250; Peter Davies), Alexa Fluor 488 goat anti-guinea pig secondary antibody (1:500; Invitrogen), Alexa Fluor 647 goat anti-rabbit IgG secondary antibody (1:500; Invitrogen), and biotinylated horse anti-mouse secondary antibody (1:200; Vector).

### Human Brain Tissue Microscopy and Analysis

Confocal images were generated using a Nikon Ti2-E microscope equipped with a Crest X-Light-V2 spinning disk confocal (Crest Optics), Celeste Light Engine (Lumencor), Piezo stage (Mad City Labs), and a Prime 95B 25mm CMOS camera (Photometrics) using a CFI Plan Apo Lambda 60x/1.4 oil or (Plan Apo VC 100×/1.4 oil) (Nikon). Images were captured using a penta dichroic 405/488/561/640/750 (Nikon), solid-state lasers 405 nm, 477 nm, 546 nm, and 638 nm and emission filters FF01-438/24, FF01-511/20, FF01-560/25, FF01-685/40 (Semrock), Nikon Multi-band for DAPI/AmyloGlo, Alexa 488, Alexa 568, and Alexa 647, respectively. The data was captured with NIS-Elements software (v. 5.41.01 build 1709, Nikon). Whole-section tiled images were acquired with an Axioscan.Z1 slide scanner (Zeiss) at 20× magnification.

Human microglial GPC4 quantitation was performed blinded to clinical and pathological diagnosis. For each brain section tiled image, 10 random ROIs of the cortex were exported into FIJI^81^. IBA^+^ microglia were segmented using FIJI’s Trainable Weka Segmentation plugin^82^ as previously described^83^. Briefly, the WEKA segmentation classifier was trained with 16 input 8-bit images (one image from each brain) using a fast random forest classifier with the following balanced training features: Gaussian blur, Hessian, membrane projections, Sobel filter, and difference of gaussians. The model was retrained for a total of 5 training sessions and then applied to each image to generate probability maps for microglia objects. The probability maps were thresholded to create microglia image masks. The image masks were then applied to the corresponding GPC4 channels to measure microglial GPC4 mean fluorescence across all brain sections. To quantify microglial GPC4 intensity as a function of distance from amyloid plaques in Alzheimer’s brains, eight Amylo-Glo-positive amyloid plaques were randomly chosen per brain section. For each plaque, IBA1^+^ microglia were defined as “near” if they were located within 125 µm from the center of the plaque. IBA1^+^ microglia were defined as “far” if they were located > 150 µm from the edge of all plaques but within 350 µm of the index plaque. Images for this analysis were 16-bit. For quantification of amyloid plaque and neurofibrillary tangle burden, whole brain section tiled images were analyzed using Zen 3.9 (Zeiss) and performed blinded to clinical and pathological diagnosis. The overlying neocortex was manually defined, and brightness thresholding was used to delineate anti-amyloid plaque and anti-CP13-positive areas. For each section, the total area of positive signal coverage was measured in the neocortex and expressed as percentage of total area analyzed in each brain.^49^

### Mouse Brain Tissue and Immunohistochemistry

Use of all animals was approved by the UVA Institutional and Animal Care and Use Committee (IACUC). All animals used in this study were handled according to IACUC approved protocols and housed in IACUC approved vivariums at the UVA MR-4. The 5xFAD^84^ (Jackson Laboratory stock #34848), APP-SAA knock-in^85^ (stock #034711), and C57BL/6J (stock #000664) mice were originally purchased from the Jackson Laboratory. All transgenic mice used were on a congenic C57BL/6 J background.

Samples from APP-SAA and matched C57BL/6J controls were collected at 9 months of age. Samples from 5xFAD and matched C57BL/6J control mice, all female, were collected at 8 months of age. Mice were anaesthetized with isoflurane and transcardially perfused with chilled PBS followed by chilled 4% PFA. Brains were post-fixed in 4% PFA on ice for 2 hours before being transferred to 30% sucrose in PBS for cryoprotection. Frozen brains were coronally sliced at 30 µm on a cryostat (Leica CM1950).

For histological staining, free-floating sections were incubated with blocking solution (5% bovine serum albumin, 2% horse serum, 1% Triton X-100 in PBS) for two h at RT before staining. Primary and secondary antibodies were diluted in the same blocking solution. Slices were incubated in primary antibodies at 4 °C for 24 h and secondary antibodies at room temperature for 2 h. The following primary antibodies were used: rabbit anti-GPC4 (ProteinTech; 1:500); chicken anti-GFAP (Aveslabs; 1:500); and goat anti-IBA1 (Abcam; 1:300). The following secondary antibodies from Jackson ImmunoResearch were used at 1:100: anti-rabbit Cy3, anti-chicken AlexaFluor 488 (703-545-155), anti-goat Cy5 (711-175-147). After secondary antibody incubation, slices were stained with Amylo-Glo (Biosensis; TR-300-AG) according to manufacturer protocols in order to detect amyloid plaques, and then were mounted using Vectashield Plus mounting media (Vector Cat#H-1900). Slides were imaged using an Innovative Imaging Innovations (3i) spinning-disc confocal microscope equipped with a Yokogawa CSX-X1 scan head using 40x and 63x objectives. Images were captured from the dentate gyrus and frontal cortex with a 63x objective for representative images and z-stacks captured with a 20x objective used for GPC4 quantification.

Image analysis was performed using Imaris software (Oxford Instruments, ver. 10.2.0). Microglia, astrocyte, and plaque surfaces were defined using automatic thresholds. After background subtraction, average intensity of the GPC4 channel within microglia and astrocyte surfaces was recorded.

### Drosophila Immunohistochemistry and Imaging

Flies were collected for staining at 10 days post eclosion (dpe). After CO_2_ anesthesia, heads were removed and fixed in 4% formaldehyde (Thermo Scientific) for 15 minutes at room temperature. Brains were dissected and incubated with primary antibodies dissolved in 0.3% Triton-X100 in PBS (PTX) for 48 h at 4 °C, followed by secondary antibodies in 0.3% PTX for 48h at 4 °C. The following primary antibodies were used: chicken anti-GFP (Aves Labs, 1:1000); rat anti-mCherry (Invitrogen, 1:1000); mouse anti-amyloid (1:200, BioLegend), and rabbit anti-cDCP1 (Cell Signaling, 1:400). The following secondary antibodies from Jackson ImmunoResearch were used at 1:100: anti-rat Cy3, anti-chicken AlexaFluor 488, and anti-rabbit Cy5. After secondary antibody incubation, slices were stained with Amylo-Glo (Biosensis; TR-300-AG) according to manufacturer protocols, and then were mounted using Vectashield Plus mounting media (Vector; H-1900). Slides were imaged using an Innovative Imaging Innovations (3i) spinning-disc confocal microscope equipped with a Yokogawa CSX-X1 scan head using 40x and 63x objectives.

### Drosophila Stocks

The following previously made *D. melanogaster* transgenes were used in this study: wrapper-Gal4DBD, Nrv2-VP16AD (CtxGlia-SplitGal4)^86^, alrm-Gal4^87^, nSyb-QF2^88^, UAS-lacZ-NLS^89^, QUAS-R-GECO1^90^, and stocks acquired from Bloomington Drosophila Stock Center (BDSC): UAS-dlpRNAi #1 (BDSC #34091), UAS-dlpRNAi #2 (BDSC #50540^91^), UAS-dlp (BDSC #9160), UAS-hGPC4 (BDSC #78408), QUAS-Aβ42 (BDSC #83347^41^).

### Drosophila Lifespan and Climbing

*D. melanogaster* crosses were set on Molasses Formula Food (Archon Scientific) at 25°C. Animals of the desired genotypes were collected at 0dpe and housed in vials containing corn syrup/soy medium (Archon Scientific) with 4-10 flies of the same sex per vial. The flies were assessed daily for viability, recorded, and remaining live flies were transferred to fresh vials every 2 days. Additionally, animals were tested for locomotion via climbing assay at 1 and 5dpe. Flies were transferred to empty 9 cm tall vials and allowed to habituate for 3 min. Vials were placed in a 9-vial holder and firmly tapped three times to knock all flies to the bottom of the vial. They were then rested for 30 s, tapped, rested again for 30 s, and received a final tap followed by the climbing assessment in which the highest height climbed by each fly in 10 s was recorded in cm. Each assay was performed between Zeitgeber time 8-9 and recorded on video. Analyses were performed using R, with lifespan analyzed with a Cox proportional hazard model with Bonferroni corrections and climbing data analyzed by a Tobit regression with Bonferroni corrections to account for floor and ceiling effects of the vial constraints.

### Phagocytosis Assays

Microglia were plated at 50% confluence on Matrigel-coated 96-well plates. All phagocytosis assays were performed in quadruplicate using pHrodo-Red-labeled substrates. FL tau fibrils or magnetic beads were added to day 8 iTF-Microglia at a final concentration of 100 nM or 1.92 × 10^7^ particles/mL, respectively. Phagocytosis was monitored every three hours for a duration of 24 h using an Incucyte SX5 (Sartorius). Four fields of view at 20x magnification per well were captured for each condition using phase and fluorescence channels. Incucyte 2023A software equipped with the Cell-by-Cell module (Sartorius) was used to create image masks of both individual cells as well as phagocytosed substrates. Cellular integrated fluorescence intensity values of each well were averaged across treatment conditions and graphed as a function of time using Prism 10 (GraphPad). Where indicated, actin polymerization was inhibited by pretreating cells with 5 μM Cytochalasin D (Millipore Sigma; C8273) for 30 min before the addition of phagocytic substrates.

### Flow Cytometry

iTF Microglia were differentiated at 15,000 cells per well in a 96-well plate. On day 8, they were treated with 1 µM Aβ_40_ or Aβ_42_ fibrils for 24 h. Cells were harvested with Versene for 8 min and resuspended and washed in DPBS plus 1% FBS, 1 mM EDTA, and 0.1% sodium azide. Immunostaining for cell-surface proteins was performed at 4 C° for 1 h with α-10E4 (Amsbio,1:200) α-GPC4 (ProteinTech; 1:200), and α-GPC6 (Bioss; 1:200) antibodies followed by secondary staining at 4 C° for 1 h with Alexa Fluor 488 goat anti-mouse IgM or Alexa Fluor 647 goat anti-rabbit IgG (1:1000). For tau fibril cell-surface binding experiments, cells were pretreated with 100 nM of anti-GPC4 sdAb-Fc for 24 h. Cells were then equilibrated for 4 °C for 15 min before the addition of tau fibrils-AF 647 for 60 min at 4 °C, washed with cold PBS, dissociated with Versene, and subjected to flow cytometry. Single cell fluorescence was measured using a Beckman Coulter CytoFlex flow cytometer. Median fluorescence intensity values of stained cells were normalized to vehicle-treated control samples and data were plotted using Prism (GraphPad, v10).

For FRET flow cytometry assays, iPSC neuron biosensors were harvested with a 1:1 mix of papain (Worthington; LK003178) and Accutase (VWR / ICT; AT-104) for 10 minutes prior to resuspending in complete iNeuron media without phenol red. FRET flow cytometry was performed on a FACSCelesta (BD Biosciences). To measure mClover and FRET, cells were excited with the 488 nm laser, and fluorescence was captured with a 530/30 nm and 610/20 nm filter, respectively. To measure mApple, cells were excited with a 561 nm laser and fluorescence was captured with a 610/20 nm filter. To quantify FRET, we used a gating strategy similar to that previously described.^49,50^ The percentage of FRET-positive cells was used for all analyses. For each experiment, 10,000 cells per replicate were analyzed and each condition was analyzed in quadruplicate. Flow cytometry data were analyzed using FlowJo (v.10.9) and data were plotted using Prism (GraphPad, v10).

### GPC4 Shedding Assays

Conditioned media samples were harvested 24 h after Aβ_40/42_ fibril treatment and clarified by centrifugation (500 x g 5 min). Cells were lifted with Versene and pelleted by centrifugation (1000 x g 5 min). The pellet was resuspended in 100 µL of Luciferase assay buffer (Promega; N112A) to lyse and generate a 1× cell lysate sample. NanoLuc Furimazine substrate (Promega; N113A) was diluted 1:10,000 substrate to assay buffer and 1:4 in PBS. To minimize potential cross-luminescence, 20 µL of each sample was loaded into 384-well assay plate (Greiner) surrounded by empty wells. 60 µL of diluted Furimazine substrate was injected and luminescence of each well was read in a Tecan M1000 Pro. The average luminescence values of conditioned media or cell lysate samples were calculated and normalized to PBS-treated samples as a control.

### Protein Pull downs

Tau-biotin (1 µg) was mixed with 3 µg of mammalian-derived GPC4 WT (Acro), 3 µg of mammalian-derived lipidated APOE3, or both in a total of 50 µL of 100 mM NaCl, 10 mM HEPES containing 0.05% Tween20. The proteins were incubated O/N at 4 °C while mixing at 700 RPM. The next day, the protein complexes were added to 50 µL of washed streptavidin Magnesphere Paramagnetic Beads (Promega; Z5482) and incubated for 1 h at RT with mixing at 700 RPM. The magnetic beads were then washed three times with TBS-T containing 0.5 M NaCl and eluted in 10 µL of 1 M glycine, pH 2 for 10 minutes at RT.

### Biolayer Interferometry

We characterized the interactions of tau with GPC4 and APOE3 via biolayer interferometry (Octet RED384, FortéBio). Assay buffer was composed of 10 mM HEPES, pH 7.4, 0.1% BSA and 0.01% Tween 20 in deionized water. Biotinylated tau-441 (rPeptide) was diluted with assay buffer to a concentration of 100 nM and Glypican 4 and APOE3 was diluted to a final concentration of 250 nM. In addition, a control anti-tau antibody, MD3.1^92^, was diluted in assay to a final concentration of 50 nM. Streptavidin biosensors (FortéBio) were equilibrated in 200 µL of assay buffer for 30 minutes before use in a 96-well plate (Greiner). For experimental analysis, 80 µL of tau-biotin, Glypican 4, APOE3, and MD3.1 were dispensed into respective wells on a 384-well assay plate. The sensors were loaded with tau-biotin, followed by quenching of the remaining streptavidin on the sensors with 10 µM biotin buffer (assay buffer with 50 mM biotin), an association step (1800 s) and a dissociation step (800 s). The assay was performed at 25 °C with 1000 rpm shaking. Data were processed and analyzed using the Octet 12.2.2.4 data analysis software. Graphs were subsequently generated in GraphPad Prism.

### Negative Stain Transmission Electron Microscopy

Fibrillization reactions were examined for the presence of filaments by negative staining and transmission electron microscopy (TEM). Briefly, 5 µL of sample was spotted on a glow-discharged carbon/formvar-coated 300-mesh grid for 1 minute. Excess liquid was blotted off with Whatman paper, followed by washing with another 5 µL of ddH_2_O. Excess liquid was removed by blotting and the grid was stained with a 2% uranyl acetate solution for 1 minute. Excess liquid was blotted off and the grid was imaged by TEM on an FEI Tecnai G2 Spirit Biotwin operating at 120 kV.

### qPCR

RNA was extracted and purified using Tri-isolate RNA Pure kits (IBI Scientific; IB47632) according to manufacturer guidelines. For qPCR assays, RNA was DNAse-treated and converted to cDNA using Quantitect Reverse-Transcription kit (Qiagen; 205311). qPCR reactions were performed using SYBR Select Master Mix (Thermo Scientific; 4472908). Final primer concentrations were 250 nM and Tm was 60 °C. Fluorescent emissions were detected using Bio-Rad CFX Connect qPCR instrument. Data were analyzed using ΔΔCT method. For representation of the data in figures, a Z-score was computed and heatmaps were generated with heatmapper.ca. qPCR primers are provided in Table 2.

### Lentiviral transduction of iPSCs with sgRNA constructs

Pooled sgRNAs were lentivirally packaged in HEK293T cells (ATCC; CRL-3216) as previously described^73^ using TransIT Lenti Reagent (Mirus Bio; MIR 6600), and introduced into CRISPRi or CRISPRa iPSCs. Cells were selected with 1 µg/mL puromycin (Gibco; A11138-03) for 7 d after which cells were cultured for 2–4 d in the absence of puromycin to allow them to recover. sgRNA protospacer sequences are provided in Table 2.

### Statistics

All cell culture data are expressed as mean ± S.E.M. from 3 or more independent experiments, and the level of significance between 2 groups was assessed with Student’s t-test. For experiments consisting of more than 2 groups, one-way or two-way ANOVA followed by Holm-Sidak test was applied. A value of p < 0.05 was considered to be statistically significant.

### Antibodies

Anti-10E4 mouse primary antibody (Amsbio; F58-10E4)

Anti-Amyloid mouse primary antibody (Millipore Sigma; MAB5206)

Anti-Amyloid mouse primary antibody (BioLegend; SIG-39320)

Anti-cleaved Dcp-1 antibody (Cell Signaling Technology; 9578)

Anti-Glypican 1 rabbit primary antibody (Abcam; EPR19285)

Anti-Glypican 2 rabbit primary antibody (Invitrogen; PA5-115299)

Anti-Glypican 3 mouse primary antibody (Invitrogen; MA5-17083)

Anti-Glypican 4 rabbit primary antibody (ProteinTech; 13048-1-AP)

Anti-Glypican 5 mouse primary antibody (R&D Systems; 297716)

Anti-Glypican 6 rabbit primary antibody (Bioss Antibodies; bs-2177R)

Anti-IBA1 rabbit primary antibody (Cell Signaling Technology; 17198)

Anti-IBA1 guinea pig primary antibody (Synaptic Systems; 234 308)

Anti-PU.1 rabbit primary antibody (Cell Signaling Technology; 2266)

Anti-Tau mouse primary antibody (CP13; gift from Peter Davies)

Anti-TMEM119 mouse primary antibody (BioLegend; 853301)

Alexa Fluor 488 donkey anti-chicken IgG secondary antibody (Jackson ImmunoResearch; 703-545-155)

Alexa Fluor 488 goat anti-guinea pig IgG secondary antibody (Invitrogen; A-11073)

Alexa Fluor 488 Plus goat anti-mouse IgG secondary antibody (Invitrogen; A32723)

Alexa Fluor 488 goat anti-mouse IgM secondary antibody (Invitrogen; A-21042)

Alexa Fluor 546 goat anti-rabbit IgG secondary antibody (Invitrogen; A-11035)

Alexa Fluor 647 goat anti-rabbit IgG secondary antibody (Invitrogen; A-21245)

Biotinylated horse anti-mouse IgG secondary antibody (Vector; BA-2000-1.5)

Cy3 donkey anti-rabbit IgG secondary antibody (Jackson ImmunoResearch; 711-165-152)

Cy3 donkey anti-rat IgG secondary antibody (Jackson ImmunoResearch; 712-165-153)

Cy5 donkey anti-goat IgG secondary antibody (Jackson ImmunoResearch; 705-175-147)

Cy5 donkey anti-rabbit IgG secondary antibody (Jackson ImmunoResearch; 711-175-152)

## Acknowledgements

This work was supported by the donors of the Alzheimer’s Disease Research Program, a program of BrightFocus Foundation (to B.B.H.), Shenandoah Foundation (to B.B.H.), UVA Brain Institute and Strategic Investment Fund (to J.C.C.), National Institutes of Health Grants 1K08NS133290 (to B.B.H.), R35GM122451 (to J.A.W.), R01NS121101 (to J.C.C.), RF1AG061874 (to C.C.), K24AG053435 (to L.T.G), and Alzheimer’s Association Zenith Fellows Award ZEN-22-969903 (to M.K.). The UCSF Neurodegenerative Disease Brain Bank is supported by NIH grants P01AG019724, P30AG062422, U19AG063911, and U01AG057195; the Rainwater Charitable Foundation, and the Bluefield Project to Cure FTD.

Data for this study were acquired at the Center for Advanced Light Microscopy at UCSF on the CREST/C2 Confocal obtained using grants from the UCSF Program for Breakthrough Biomedical Research funded in part by the Sandler Foundation and the UCSF Research Resource Fund Award. We thank John Lukens, PhD, Lulu Jiang, MD PhD, and Marc Diamond, MD for supplying reagents. We thank Irene Lui, PhD, for technical assistance with mass spectrometry.

## REFERENCES

1. Braak, H., Alafuzoff, I., Arzberger, T., Kretzschmar, H. & Del Tredici, K. Staging of Alzheimer disease-associated neurofibrillary pathology using paraffin sections and immunocytochemistry. Acta Neuropathol 112, 389–404 (2006).

2. Musiek, E. S. & Holtzman, D. M. Origins of Alzheimer’s disease: reconciling cerebrospinal fluid biomarker and neuropathology data regarding the temporal sequence of amyloid-beta and tau involvement. Current Opinion in Neurology **Publish Ahead of Print**, 10.1097/WCO.0b013e32835a30f4 (2012).

3. Götz, J., Chen, F., van Dorpe, J. & Nitsch, R. M. Formation of Neurofibrillary Tangles in P301L Tau Transgenic Mice Induced by Aβ42 Fibrils. Science 293, 1491–1495 (2001).

4. Lewis, J. et al. Enhanced neurofibrillary degeneration in transgenic mice expressing mutant tau and APP. Science 293, 1487–1491 (2001).

5. Hurtado, D. E. et al. A{beta} accelerates the spatiotemporal progression of tau pathology and augments tau amyloidosis in an Alzheimer mouse model. Am J Pathol 177, 1977–1988 (2010).

6. Jagust, W. Imaging the evolution and pathophysiology of Alzheimer disease. Nat Rev Neurosci 19, 687–700 (2018).

7. Vogel, J. W. et al. Spread of pathological tau proteins through communicating neurons in human Alzheimer’s disease. Nat Commun 11, 2612 (2020).

8. Lee, W. J. et al. Regional Aβ-tau interactions promote onset and acceleration of Alzheimer’s disease tau spreading. Neuron 110, 1932–1943.e5 (2022).

9. Van Dyck, C. H. et al. Lecanemab in Early Alzheimer’s Disease. N Engl J Med 388, 9–21 (2023).

10. Knopman, D. S. Lecanemab reduces brain amyloid-β and delays cognitive worsening. Cell Rep Med 4, 100982 (2023).

11. Shcherbinin, S. et al. Association of Amyloid Reduction After Donanemab Treatment With Tau Pathology and Clinical Outcomes: The TRAILBLAZER-ALZ Randomized Clinical Trial. JAMA Neurology 79, 1015–1024 (2022).

12. Naj, A. C. et al. Common variants at MS4A4/MS4A6E, CD2AP, CD33 and EPHA1 are associated with late-onset Alzheimer’s disease. Nat Genet 43, 436–41 (2011).

13. Zhang, Y. et al. An RNA-sequencing transcriptome and splicing database of glia, neurons, and vascular cells of the cerebral cortex. J Neurosci 34, 11929–11947 (2014).

14. Nott, A. et al. Brain cell type-specific enhancer-promoter interactome maps and disease risk association. Science (2019) doi:10.1126/science.aay0793.

15. Novikova, G. et al. Integration of Alzheimer’s disease genetics and myeloid genomics identifies disease risk regulatory elements and genes. Nature Communications 12, 1610 (2021).

16. Zhang, B. et al. Integrated systems approach identifies genetic nodes and networks in late-onset Alzheimer’s disease. Cell 153, 707–20 (2013).

17. Podleśny-Drabiniok, A., Marcora, E. & Goate, A. M. Microglial Phagocytosis: A Disease-Associated Process Emerging from Alzheimer’s Disease Genetics. Trends Neurosci (2020) doi:10.1016/j.tins.2020.10.002.

18. Bhaskar, K. et al. Regulation of tau pathology by the microglial fractalkine receptor. Neuron 68, 19–31 (2010).

19. Asai, H. et al. Depletion of microglia and inhibition of exosome synthesis halt tau propagation. Nat Neurosci 18, 1584–1593 (2015).

20. Maphis, N. et al. Reactive microglia drive tau pathology and contribute to the spreading of pathological tau in the brain. Brain 138, 1738–1755 (2015).

21. Johnson, N. R. et al. CSF1R inhibitors induce a sex-specific resilient microglial phenotype and functional rescue in a tauopathy mouse model. Nat Commun 14, 118 (2023).

22. Shi, Y. et al. ApoE4 markedly exacerbates tau-mediated neurodegeneration in a mouse model of tauopathy. Nature 549, 523–527 (2017).

23. Shi, Y. et al. Microglia drive APOE-dependent neurodegeneration in a tauopathy mouse model. J Exp Med 216, 2546–2561 (2019).

24. Pascoal, T. A. et al. Microglial activation and tau propagate jointly across Braak stages. Nat Med 27, 1592–1599 (2021).

25. Eikelenboom, P. et al. Neuroinflammation - an early event in both the history and pathogenesis of Alzheimer’s disease. Neurodegener Dis 7, 38–41 (2010).

26. Yoshiyama, Y. et al. Synapse loss and microglial activation precede tangles in a P301S tauopathy mouse model. Neuron 53, 337–51 (2007).

27. Kirkemo, L. L. et al. Cell-surface tethered promiscuous biotinylators enable comparative small-scale surface proteomic analysis of human extracellular vesicles and cells. eLife 11, e73982 (2022).

28. Keren-Shaul, H. et al. A Unique Microglia Type Associated with Restricting Development of Alzheimer’s Disease. Cell 169, 1276–1290 e17 (2017).

29. Deczkowska, A. et al. Disease-Associated Microglia: A Universal Immune Sensor of Neurodegeneration. Cell 173, 1073–1081 (2018).

30. Nimmerjahn, A., Kirchhoff, F. & Helmchen, F. Resting Microglial Cells Are Highly Dynamic Surveillants of Brain Parenchyma in Vivo. Science 308, 1314–1318 (2005).

31. Hickman, S. E. et al. The microglial sensome revealed by direct RNA sequencing. Nature neuroscience 16, 1896–905 (2013).

32. Dräger, N. M. et al. A CRISPRi/a platform in human iPSC-derived microglia uncovers regulators of disease states. Nat Neurosci (2022) doi:10.1038/s41593-022-01131-4.

33. Snow, A. D. et al. The presence of heparan sulfate proteoglycans in the neuritic plaques and congophilic angiopathy in Alzheimer’s disease. Am J Pathol 133, 456–463 (1988).

34. Su, J. H., Cummings, B. J. & Cotman, C. W. Localization of heparan sulfate glycosaminoglycan and proteoglycan core protein in aged brain and Alzheimer’s disease. Neuroscience 51, 801–813 (1992).

35. Holmes, B. B. et al. Heparan sulfate proteoglycans mediate internalization and propagation of specific proteopathic seeds. Proc Natl Acad Sci U S A 110, E3138–47 (2013).

36. Stopschinski, B. E. et al. Specific glycosaminoglycan chain length and sulfation patterns are required for cell uptake of tau vs. alpha-synuclein and beta-amyloid aggregates. The Journal of biological chemistry (2018) doi:10.1074/jbc.RA117.000378.

37. Rauch, J. N. et al. Tau Internalization is Regulated by 6-O Sulfation on Heparan Sulfate Proteoglycans (HSPGs). Sci Rep 8, 6382 (2018).

38. Schmued, L. et al. Introducing Amylo-Glo, a novel fluorescent amyloid specific histochemical tracer especially suited for multiple labeling and large scale quantification studies. Journal of Neuroscience Methods 209, 120–126 (2012).

39. Geirsdottir, L. et al. Cross-Species Single-Cell Analysis Reveals Divergence of the Primate Microglia Program. Cell 179, 1609–1622 e16 (2019).

40. Pembroke, W. G., Hartl, C. L. & Geschwind, D. H. Evolutionary conservation and divergence of the human brain transcriptome. Genome Biol 22, 52 (2021).

41. Li, Y. X., Sibon, O. C. M. & Dijkers, P. F. Inhibition of NF-κB in astrocytes is sufficient to delay neurodegeneration induced by proteotoxicity in neurons. Journal of Neuroinflammation 15, 261 (2018).

42. Stopschinski, B. E. et al. A synthetic heparinoid blocks tau aggregate cell uptake and amplification. J Biol Chem (2020) doi:10.1074/jbc.RA119.010353.

43. Dono, Rosanna. Anti-GPC4 Single Domain Antibodies. (2021).

44. Huang, K. & Park, S. Heparan Sulfated Glypican-4 Is Released from Astrocytes by Proteolytic Shedding and GPI-Anchor Cleavage Mechanisms. eNeuro 8, ENEURO.0069-21.2021 (2021).

45. Allen, N. J. et al. Astrocyte glypicans 4 and 6 promote formation of excitatory synapses via GluA1 AMPA receptors. Nature 486, 410–414 (2012).

46. Chen, Y. et al. APOE3ch alters microglial response and suppresses Aβ-induced tau seeding and spread. Cell 0, (2023).

47. Nelson, M. R. et al. The APOE-R136S mutation protects against APOE4-driven Tau pathology, neurodegeneration and neuroinflammation. Nat Neurosci 1–18 (2023) doi:10.1038/s41593-023-01480-8.

48. McQuade, A. et al. Gene expression and functional deficits underlie TREM2-knockout microglia responses in human models of Alzheimer’s disease. Nat Commun 11, 5370 (2020).

49. Holmes, B. B. et al. Proteopathic tau seeding predicts tauopathy in vivo. Proceedings of the National Academy of Sciences of the United States of America (2014) doi:10.1073/pnas.1411649111.

50. Furman, J. L., Holmes, B. B. & Diamond, M. I. Sensitive Detection of Proteopathic Seeding Activity with FRET Flow Cytometry. Journal of visualized experiments : JoVE e53205 (2015) doi:10.3791/53205.

51. Haney, M. S. et al. APOE4/4 is linked to damaging lipid droplets in Alzheimer’s disease microglia. Nature 1–8 (2024) doi:10.1038/s41586-024-07185-7.

52. Saroja, S. R., Gorbachev, K., Julia, T., Goate, A. M. & Pereira, A. C. Astrocyte-secreted glypican-4 drives APOE4-dependent tau hyperphosphorylation. Proceedings of the National Academy of Sciences 119, e2108870119 (2022).

53. Funk, K. E., Mirbaha, H., Jiang, H., Holtzman, D. M. & Diamond, M. I. Distinct Therapeutic Mechanisms of Tau Antibodies: PROMOTING MICROGLIAL CLEARANCE VERSUS BLOCKING NEURONAL UPTAKE. The Journal of biological chemistry 290, 21652–62 (2015).

54. Falkon, K. F. et al. Microglia internalize tau monomers and fibrils using distinct receptors but similar mechanisms. Alzheimers Dement (2024) doi:10.1002/alz.14418.

55. Ma, K. et al. Glypican 4 Regulates Aβ Internalization in Neural Stem Cells Partly via Low-Density Lipoprotein Receptor-Related Protein 1. Frontiers in Cellular Neuroscience 15, (2021).

56. Puliatti, G. et al. Intracellular accumulation of tau oligomers in astrocytes and their synaptotoxic action rely on Amyloid Precursor Protein Intracellular Domain-dependent expression of Glypican-4. Prog Neurobiol 102482 (2023) doi:10.1016/j.pneurobio.2023.102482.

57. de Wit, J. et al. Unbiased discovery of glypican as a receptor for LRRTM4 in regulating excitatory synapse development. Neuron 79, 696–711 (2013).

58. Hagihara, K., Watanabe, K., Chun, J. & Yamaguchi, Y. Glypican-4 is an FGF2-binding heparan sulfate proteoglycan expressed in neural precursor cells. Dev Dyn 219, 353–67 (2000).

59. Stapornwongkul, K. S., de Gennes, M., Cocconi, L., Salbreux, G. & Vincent, J.-P. Patterning and growth control in vivo by an engineered GFP gradient. Science 370, 321–327 (2020).

60. McGough, I. J. et al. Glypicans shield the Wnt lipid moiety to enable signalling at a distance. Nature 585, 85–90 (2020).

61. Raj, A., Kuceyeski, A. & Weiner, M. A Network Diffusion Model of Disease Progression in Dementia. Neuron 73, 1204–1215 (2012).

62. Zhou, J., Gennatas, E. D., Kramer, J. H., Miller, B. L. & Seeley, W. W. Predicting Regional Neurodegeneration from the Healthy Brain Functional Connectome. Neuron 73, 1216–1227 (2012).

63. Mahley, R. W. Heparan sulfate proteoglycan/low density lipoprotein receptor-related protein pathway involved in type III hyperlipoproteinemia and Alzheimer’s disease. Isr J Med Sci 32, 414–429 (1996).

64. Chen, G. et al. ApoE3 R136S binds to Tau and blocks its propagation, suppressing neurodegeneration in mice with Alzheimer’s disease. Neuron S0896-6273(24)00914–0 (2025) doi:10.1016/j.neuron.2024.12.015.

65. Grubman, A. et al. A single-cell atlas of entorhinal cortex from individuals with Alzheimer’s disease reveals cell-type-specific gene expression regulation. Nat Neurosci 22, 2087–2097 (2019).

66. Mathys, H. et al. Single-cell transcriptomic analysis of Alzheimer’s disease. Nature 570, 332–337 (2019).

67. Olah, M. et al. Single cell RNA sequencing of human microglia uncovers a subset associated with Alzheimer’s disease. Nat Commun 11, 6129 (2020).

68. Hopp, S. C. et al. The role of microglia in processing and spreading of bioactive tau seeds in Alzheimer’s disease. J Neuroinflammation 15, 269 (2018).

69. Zhu, B. et al. Trem2 deletion enhances tau dispersion and pathology through microglia exosomes. Mol Neurodegener 17, 58 (2022).

70. Fowler, S. L. et al. Tau filaments are tethered within brain extracellular vesicles in Alzheimer’s disease. Nat Neurosci 1–9 (2024) doi:10.1038/s41593-024-01801-5.

71. Libeu, C. P. et al. New Insights into the Heparan Sulfate Proteoglycan-binding Activity of Apolipoprotein E*. Journal of Biological Chemistry 276, 39138–39144 (2001).

72. Mah, D., et al. Apolipoprotein E Recognizes Alzheimer’s Disease Associated 3-O Sulfation of Heparan Sulfate. Angewandte Chemie International Edition 62, e202212636 (2023).

73. Tian, R. et al. CRISPR Interference-Based Platform for Multimodal Genetic Screens in Human iPSC-Derived Neurons. Neuron 104, 239–255.e12 (2019).

74. Leng, K. et al. CRISPRi screens in human iPSC-derived astrocytes elucidate regulators of distinct inflammatory reactive states. Nat Neurosci 25, 1528–1542 (2022).

75. Chen, D. et al. FTD-tau S320F mutation stabilizes local structure and allosterically promotes amyloid motif-dependent aggregation. Nat Commun 14, 1625 (2023).

76. Montine, T. J. et al. National Institute on Aging-Alzheimer’s Association guidelines for the neuropathologic assessment of Alzheimer’s disease: a practical approach. Acta Neuropathol 123, 1–11 (2012).

77. Alafuzoff, I. et al. Staging of neurofibrillary pathology in Alzheimer’s disease: a study of the BrainNet Europe Consortium. Brain Pathol 18, 484–496 (2008).

78. Alafuzoff, I. et al. Assessment of beta-amyloid deposits in human brain: a study of the BrainNet Europe Consortium. Acta Neuropathol 117, 309–320 (2009).

79. Cairns, N. J. et al. Neuropathologic diagnostic and nosologic criteria for frontotemporal lobar degeneration: consensus of the Consortium for Frontotemporal Lobar Degeneration. Acta Neuropathol 114, 5–22 (2007).

80. Duong, H. & Han, M. A multispectral LED array for the reduction of background autofluorescence in brain tissue. J Neurosci Methods 220, 46–54 (2013).

81. Schindelin, J., et al. Fiji: an open-source platform for biological-image analysis. Nat Methods 9, 676–682 (2012).

82. Arganda-Carreras, I. et al. Trainable Weka Segmentation: a machine learning tool for microscopy pixel classification. Bioinformatics 33, 2424–2426 (2017).

83. Kataras, T. J. et al. ACCT is a fast and accessible automatic cell counting tool using machine learning for 2D image segmentation. Sci Rep 13, 8213 (2023).

84. Oakley, H. et al. Intraneuronal β-Amyloid Aggregates, Neurodegeneration, and Neuron Loss in Transgenic Mice with Five Familial Alzheimer’s Disease Mutations: Potential Factors in Amyloid Plaque Formation. J. Neurosci. 26, 10129–10140 (2006).

85. Xia, D. et al. Novel App knock-in mouse model shows key features of amyloid pathology and reveals profound metabolic dysregulation of microglia. Molecular Neurodegeneration 17, 41 (2022).

86. Coutinho-Budd, J. C., Sheehan, A. E. & Freeman, M. R. The secreted neurotrophin Spätzle 3 promotes glial morphogenesis and supports neuronal survival and function. Genes Dev. 31, 2023–2038 (2017).

87. Doherty, J., Logan, M. A., Taşdemir, Ö. E. & Freeman, M. R. Ensheathing Glia Function as Phagocytes in the Adult Drosophila Brain. J. Neurosci. 29, 4768–4781 (2009).

88. Riabinina, O. et al. Improved and expanded Q-system reagents for genetic manipulations. Nat Methods 12, 219–222 (2015).

89. Doherty, D., Feger, G., Younger-Shepherd, S., Jan, L. Y. & Jan, Y. N. Delta is a ventral to dorsal signal complementary to Serrate, another Notch ligand, in Drosophila wing formation. Genes Dev 10, 421–434 (1996).

90. Ma, Z. & Freeman, M. R. TrpML-mediated astrocyte microdomain Ca2+ transients regulate astrocyte–tracheal interactions. eLife 9, e58952 (2020).

91. Perkins, L. A. et al. The Transgenic RNAi Project at Harvard Medical School: Resources and Validation. Genetics 201, 843–852 (2015).

92. Hitt, B. D. et al. Anti-tau antibodies targeting a conformation-dependent epitope selectively bind seeds. Journal of Biological Chemistry 299, 105252 (2023).

