## Supplemental Figures for "β-Amyloid Induces Microglial Expression of GPC4 and APOE Leading to Increased Neuronal Tau Pathology and Toxicity"

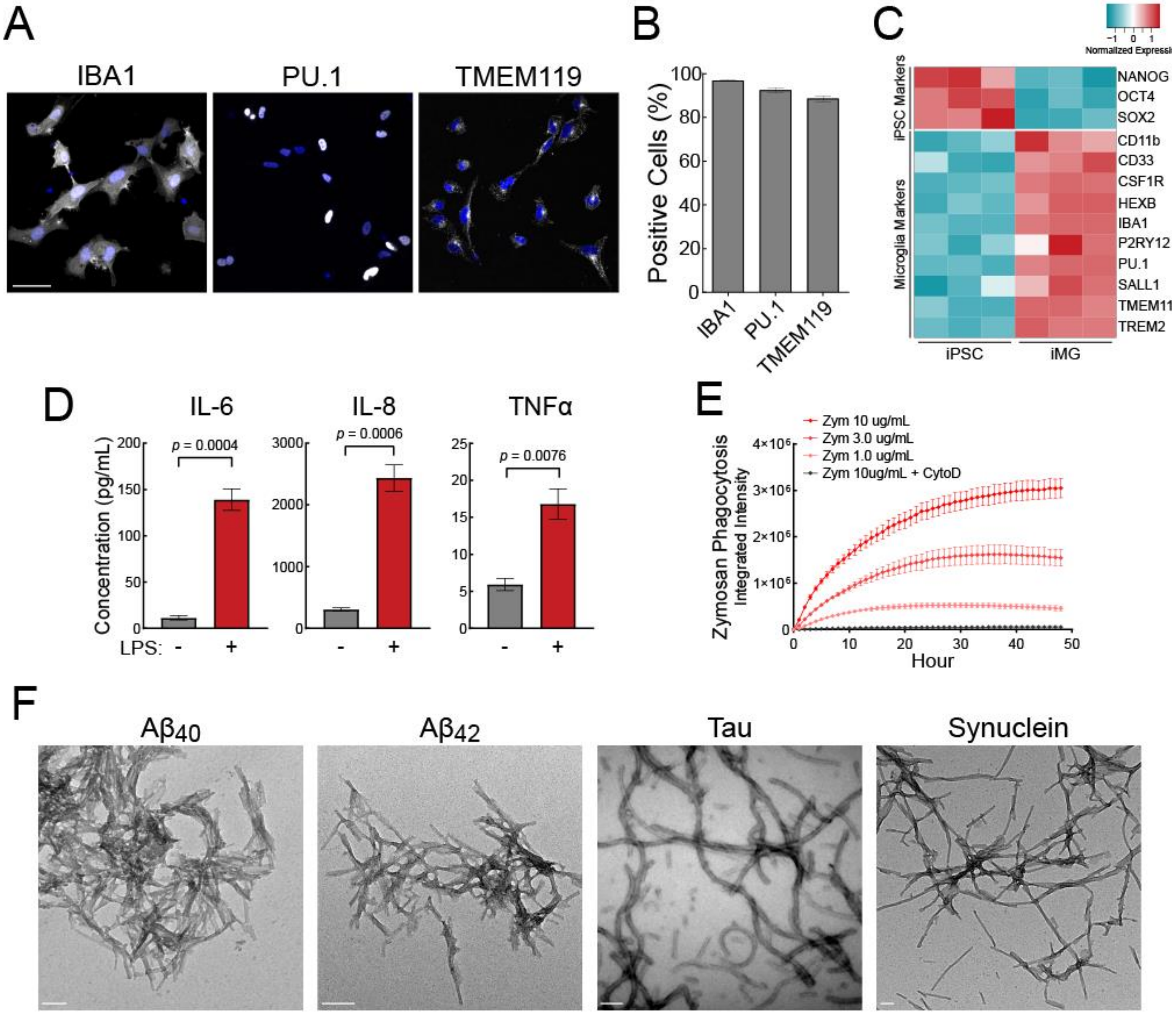


**Figure S1: Characterization of iTF Microglia and amyloids.** (A) Representative immunofluorescence micrographs of iTF-Microglia on day 8 of differentiation. iTF Microglia were stained for microglia markers IBA1, PU.1, and TMEM119. Nuclei were labeled by Hoechst 33342. Scale bar = 50 µm (B) Flow cytometry quantification of IBA1$^{+}$, PU.1$^{+}$, and TMEM119$^{+}$ cells among day-8 iTF Microglia. (C) Heat map of normalized expression of key stem cell and microglial genes in iTF iPSC or iTF Microglia cells after 0 days or 8 days of differentiation via qRT-PCR. Fold change was calculated using the ΔΔCT method with GAPDH as an endogenous control. Data was standardized to iTF-Microglia NANOG expression. (D) Laser bead immunoassay shows iTF microglia cytokine secretion of IL-6, IL-8, and TNFα in response to 100 ng/mL of lipopolysaccharide after 24 h. N = 3 biological replicates. The *p*-values were determined by Student t-test. (E) Phagocytosis of pHrodo-Red Zymosan bioparticles with and without actin polymerization inhibitor Cytochalasin D (5 µM) by iTF-Microglia. Images were taken every hour for 48 h with an IncuCyte SX5 live-cell imaging system. In all graphs, the data represent the means ± SEM. (F) Transmission electron microscopy of protein amyloids demonstrating filamentous ultrastructure. Scale bar = 100 nm.

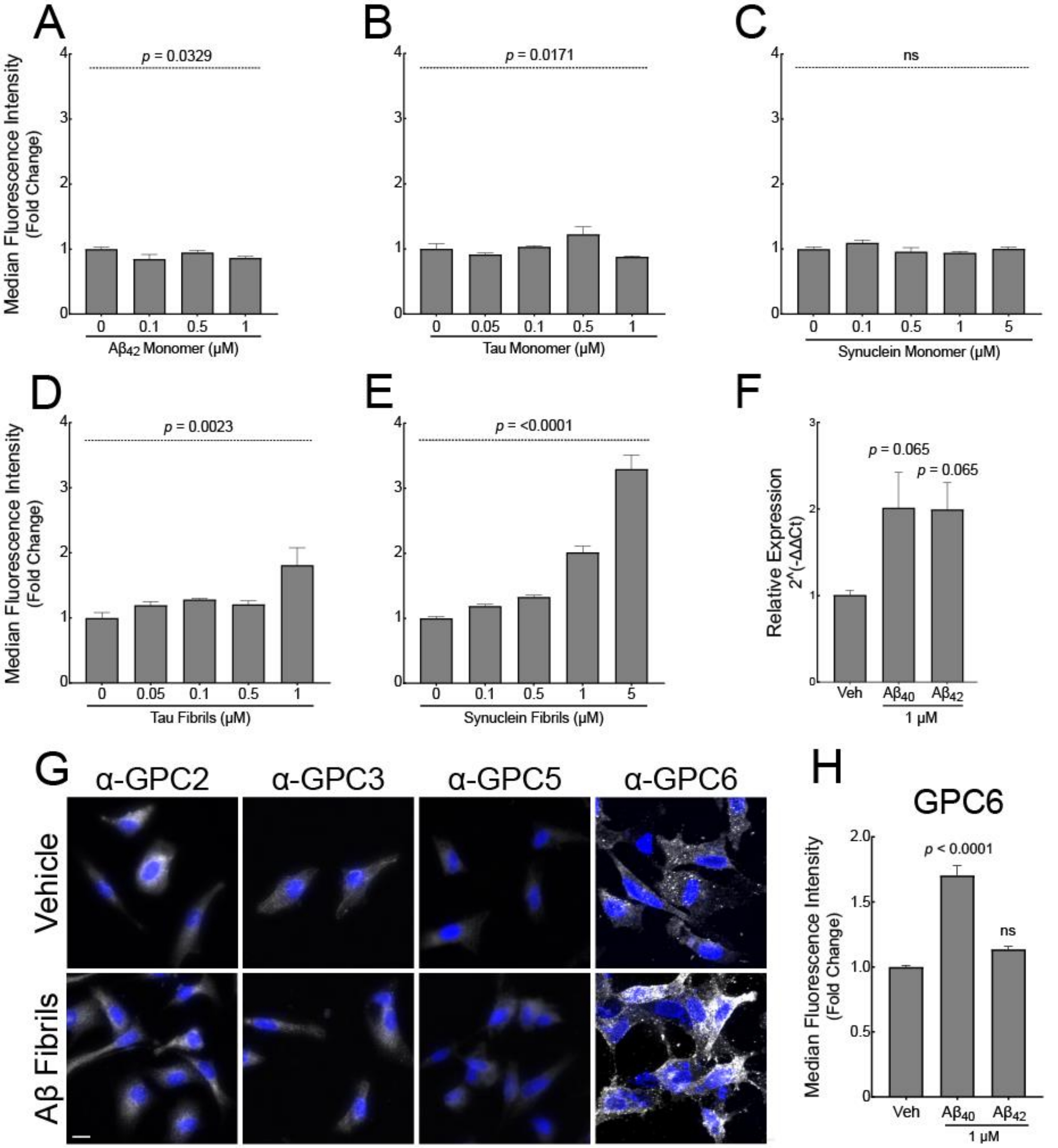


**Figure S2: Fibrils, but not monomer, induces cell-surface heparan sulfate and GPC4 expression.** iTF-Microglia treated with (A) $A\beta_{42}$, (B) tau, or (C) synuclein monomer do not increase cell-surface heparan sulfate, but cells treated with (D) tau or (E) synuclein fibrils do increase cell-surface heparan sulfate as measured by flow cytometry and 10E4 staining. The statistical analyses were performed with a one-way ANOVA. (F) Normalized GPC4 mRNA expression in iTF Microglia cells treated with amyloids and quantified by qRT-PCR. Fold change was calculated using the ΔΔCT method with GAPDH as an endogenous control. (G) GPC2, GPC3, GPC5,and GPC6 immunocytochemistry of iTF Microglia treated with 1 μM $A\beta_{40}$ for 24 h. Scale bar = 10 μm. (H) GPC6 flow cytometry quantification of iTF Microglia treated with 1 μM $A\beta_{40}$ or $A\beta_{42}$ fibrils for 24 h. The statistical analyses in F and H were performed with a one-way ANOVA and Holm-Sidak multiple comparisons test for the adjusted *p*-values. N = 3 biological replicates. In all graphs, the data represent the means ± SEM.

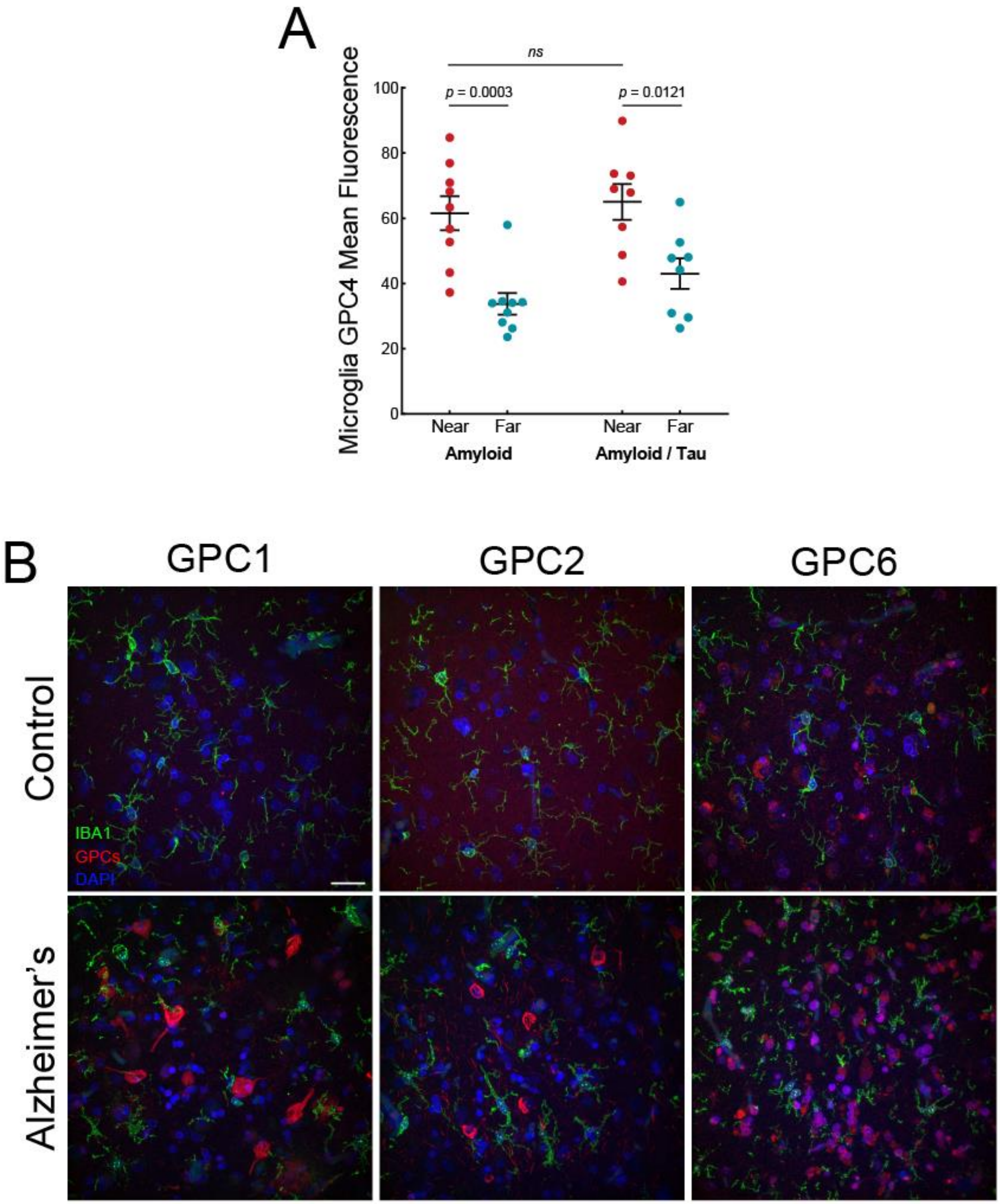


**Figure S3: Glypicans are upregulated in Human AD.** (A) Human quantification of GPC4 mean intensity values in IBA$^+$ microglia located at two distances from Amylo-Glo$^+$ Aβ plaques. A total of eight plaques were measured from two brains, and for each plaque, microglia were binned into two separate categories, "near" or "far", based on proximity to the plaque. The data compares a patient with amyloid pathology only versus a patient with amyloid and tau pathology. (B) Representative confocal images of IBA1 (green), GPCs (red), and DAPI (blue) in an age-matched control and AD case. GPC1, GPC2, and GPC6 are upregulated in AD brain but do not localize to microglia. Scale bar = 40 µm.

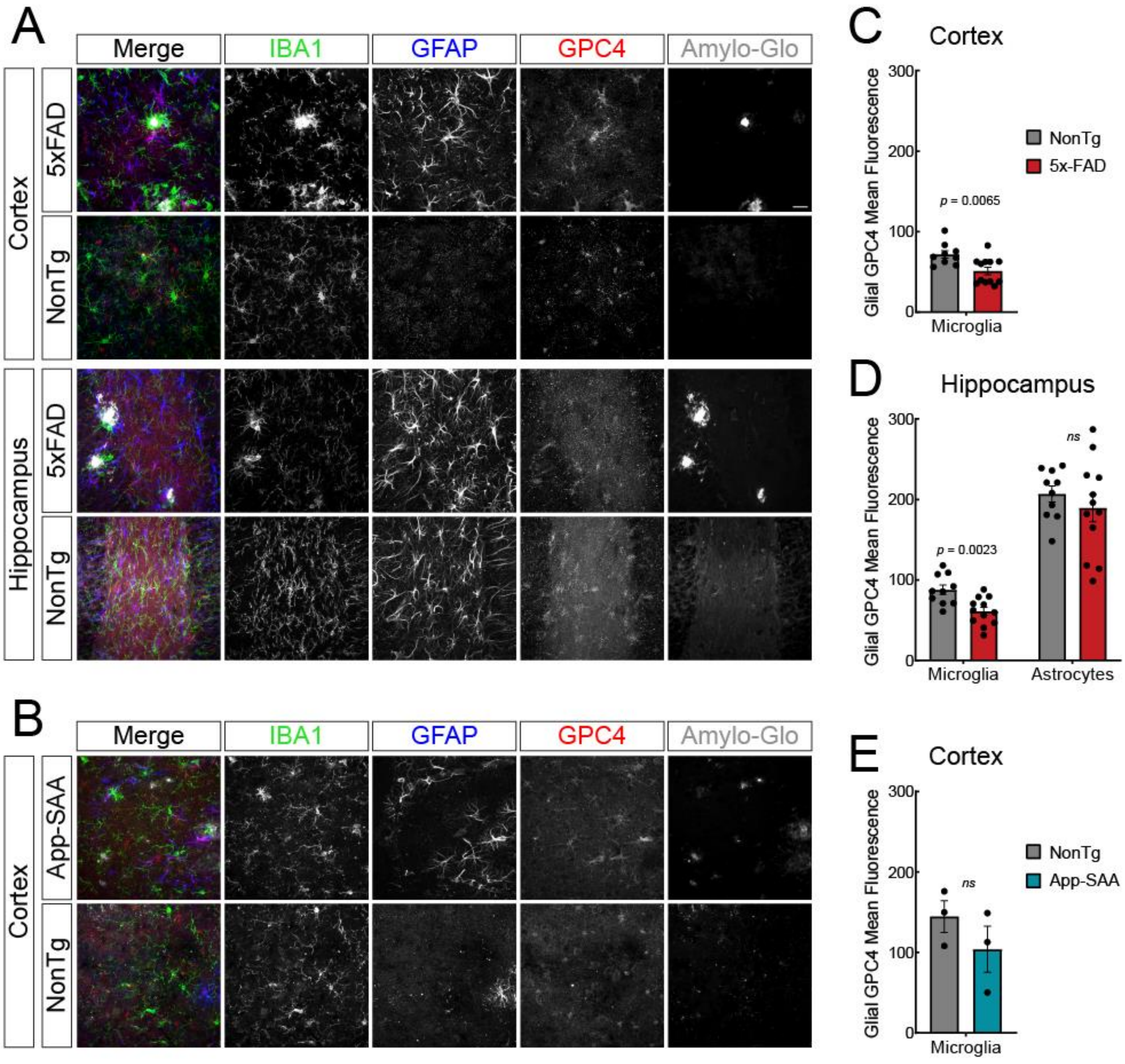


**Figure S4**: **Glia do not upregulate GPC4 in mouse models of amyloidosis.** Representative confocal images of IBA1 (green), GFAP (blue), GPC4 (red), and Amylo-Glo (white) in the cortex and hippocampus of (A) 5xFAD and (B) App-SAA mice with their corresponding age-matched non-transgenic controls. Scale bar = 10 μM. Mouse quantification of GPC4 mean intensity values in $IBA^+$ microglia or $GFAP^+$ astrocytes in the cortex or hippocampus of 5xFAD (C, D) and App-SAA mice (E). The statistical analyses were performed with a Student t-test for averaged values from individual mice. In all graphs, the data represent the means ± SEM.

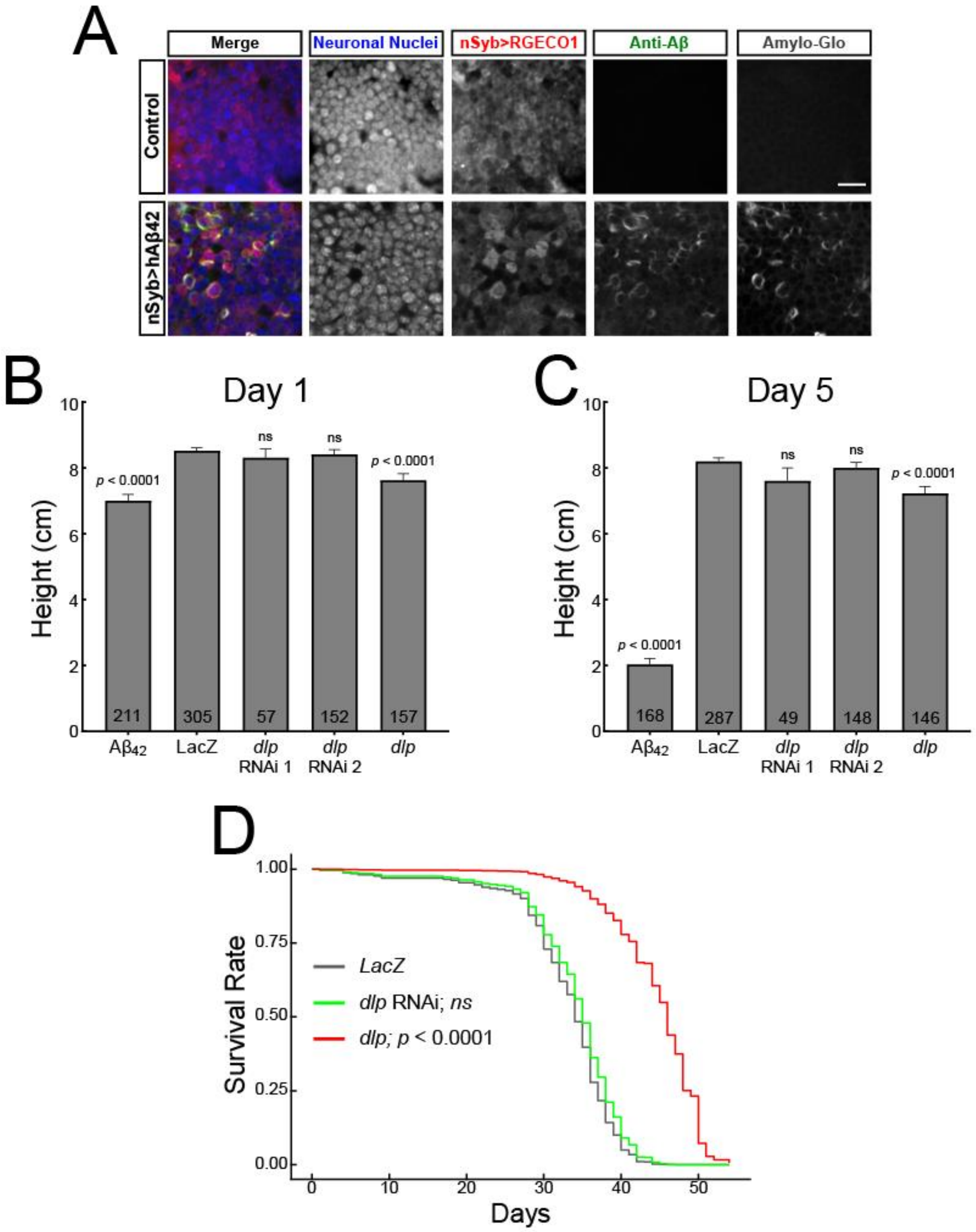


**Figure S5**: **Glial over-expression or knockdown of *dlp* minimally alters climbing or lethality in non-transgenic flies**. (A) Amyloid deposits are present within $A\beta_{42}$-expressing *Drosophila* brains. The cortex of adult control (top) and $A\beta_{42}$ (bottom) brains showing neuronal nuclei (anti-Elav, blue) and neurons expressing nSyb-QF2 (QUAS-RGECO1; red). Both total Aβ (anti-amyloid 6E10 antibody, green) and fibrillar Aβ (Amylo-Glo, white) are present in flies expressing $A\beta_{42}$ driven by nSyb-QF2 and absent in control flies. Scale bar = 10 µm. Full genotypes: (Top) w;Alrm-Gal4,wrapper-Gal4DBD,Nrv2-VP16AD,UASCD8GFP/+;nSyb-QF2,QUAS-RGECO1/+ (control) and (Bottom) w;Alrm-Gal4,wrapper-Gal4DBD,Nrv2-VP16AD,UASCD8GFP,QUAS-hAβ42/+;nSyb-QF2,QUAS-RGECO1/+ (nSyb>hAβ42). Climbing heights were measured at day 1 post-eclosion (B) or day 5 post-eclosion (C) in non-transgenic flies expressing LacZ, *dlp* RNAi, or *dlp* cDNA from a glial driver. The statistical analyses were performed with a Tobit regression with Bonferroni correction. (D) Median fly lifespans measured in days after eclosion (LacZ = 34 d; $A\beta_{42}$ = 16 d; dlp RNAi = 36 d; *dlp* = 46 d). Lifespan data was analyzed using a Cox proportional hazard model with Bonferroni corrections (LacZ n = 174; *dlp* RNAi n = 61; *dlp* n = 91). The data from the LacZ and $A\beta_{42}$ *Drosophila* lines were the same data from Figure 4.

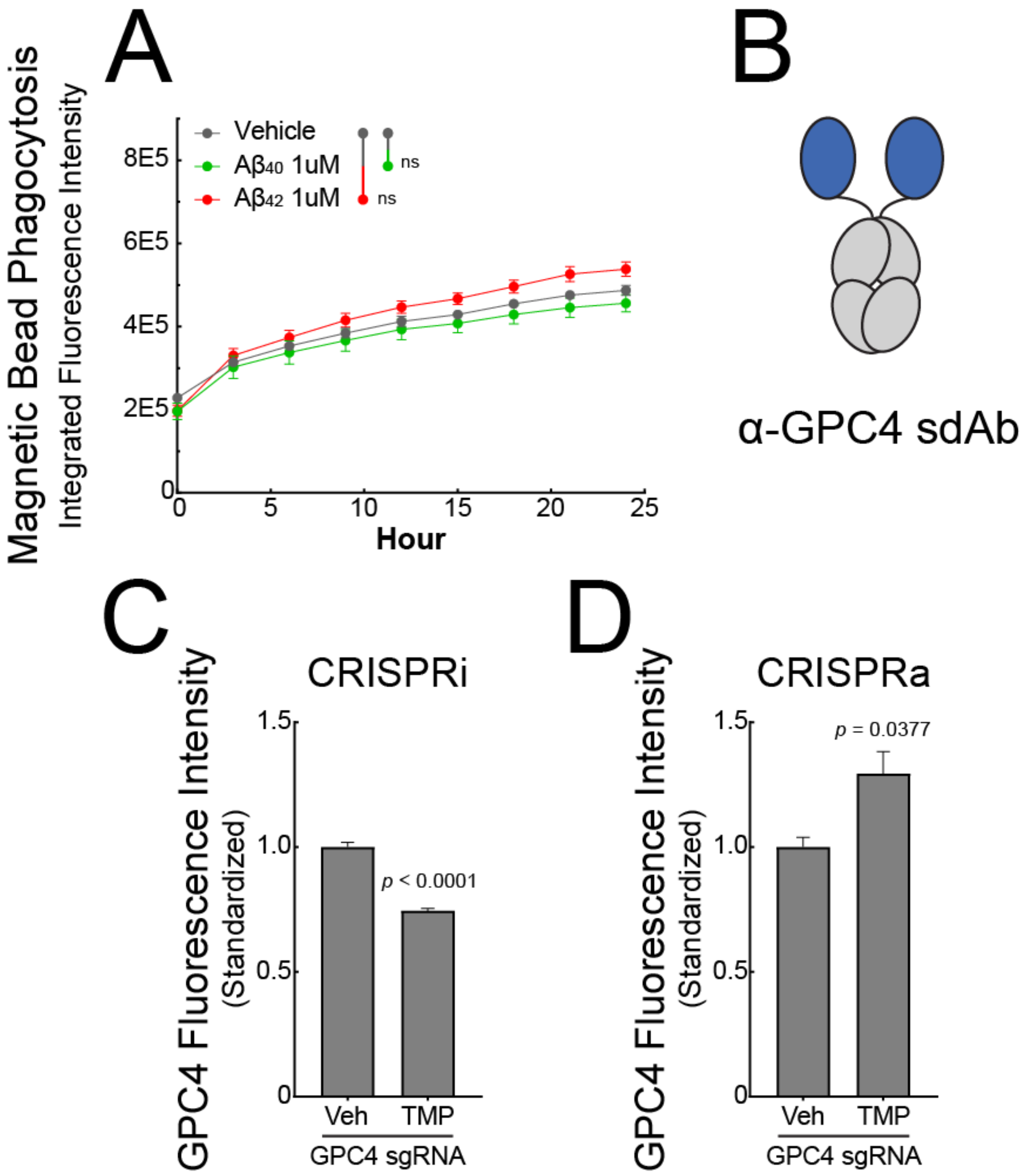


**Figure S6: HSPGs do not mediate phagocytosis of magnetic beads.** (A) Phagocytosis of pHrodo red-labeled magnetic beads by iTF-Microglia does not increase after pretreatment with $A\beta_{40}$ or $A\beta_{42}$ fibrils. (B) Cartoon depiction of engineered α-GPC4 sdAb-Fc. Cell-surface GPC4 protein levels after (C) CRISPRi or (D) CRISPRa machinery is activated by trimethoprim (TMP, 50 nM) as measured by flow cytometry. The *p*-values were determined by Student t-test. N = 3 replicates. The data represent the means ± SEM.

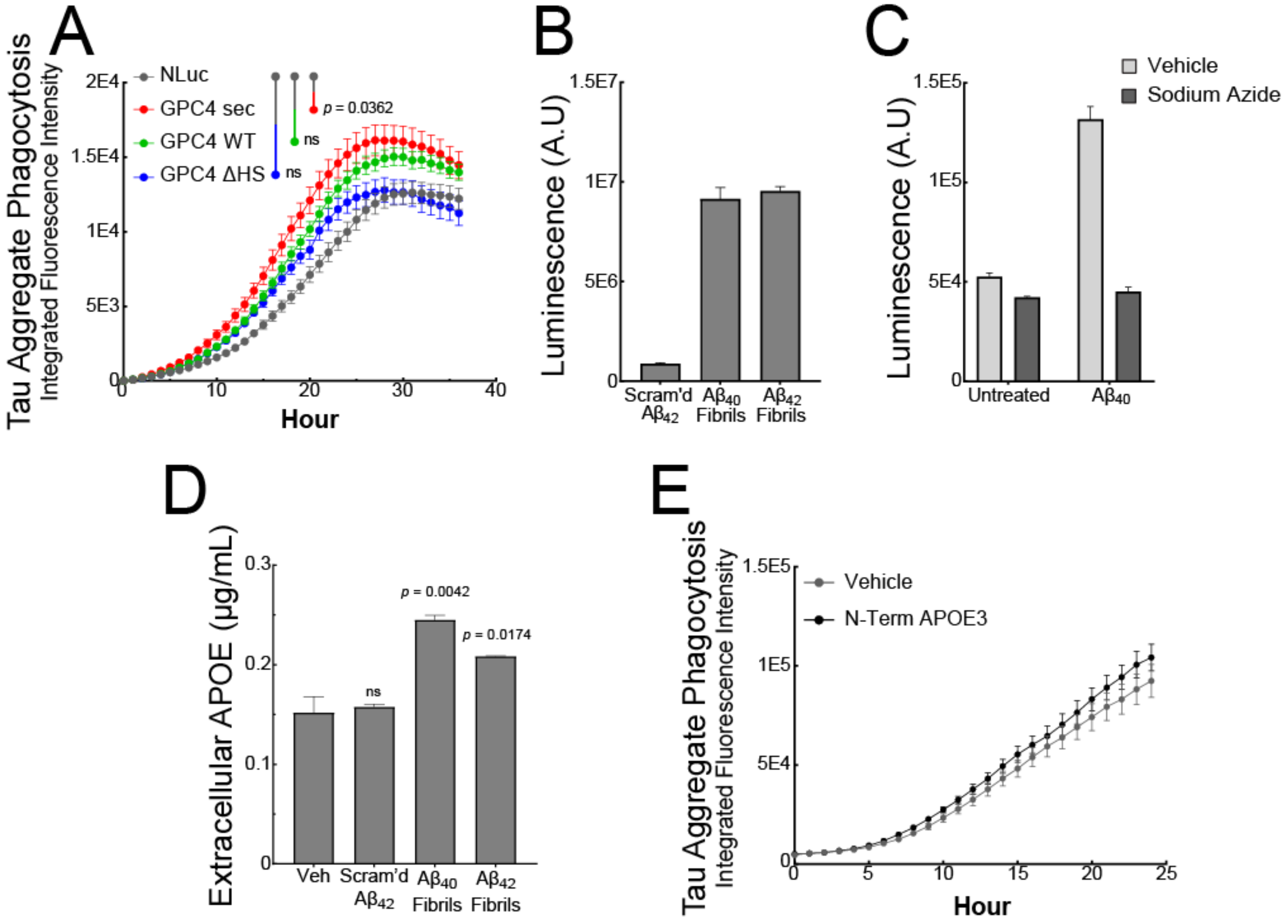


**Figure S7**: **GPC4 Shedding does not occur with scrambled A$\beta_{42}$ and requires cell metabolism.** (A) GPC4 genetic variants and a NLuc control lentivirus were transduced into BV2 cells and pHrodo-red labeled tau aggregates were measured via live-cell imaging on an Incucyte SX5. The statistical analyses were performed with a one-way ANOVA and Holm-Sidak multiple comparisons tests for the adjusted *p*-values. (B) NLuc-GPC4 luminescence was measured in iTF Microglia conditioned media after a 24 h treatment with 1 µM scrambled A$\beta_{42}$, A$\beta_{40}$ fibrils, or A$\beta_{42}$ fibrils. (C) NLuc-GPC4 luminescence was measured in iTF Microglia conditioned media after a 3 h treatment with A$\beta_{40}$ and A$\beta_{42}$ fibrils in the presence of 0.1% sodium azide. (D) iTF Microglia were treated with 1 µM scrambled A$\beta_{42}$, A$\beta_{40}$ fibrils or A$\beta_{42}$ fibrils for 24 h and the conditioned media was subjected to APOE quantification via ELISA. (E) pHrodo-red labeled tau aggregates (50 nM) were preincubated with soluble recombinant N-terminal APOE3 (250 nM) lacking the C-terminal domain and uptake was measured every hour for 24 h. N = 4 biological replicates. In all graphs, the data represent the means ± SEM.

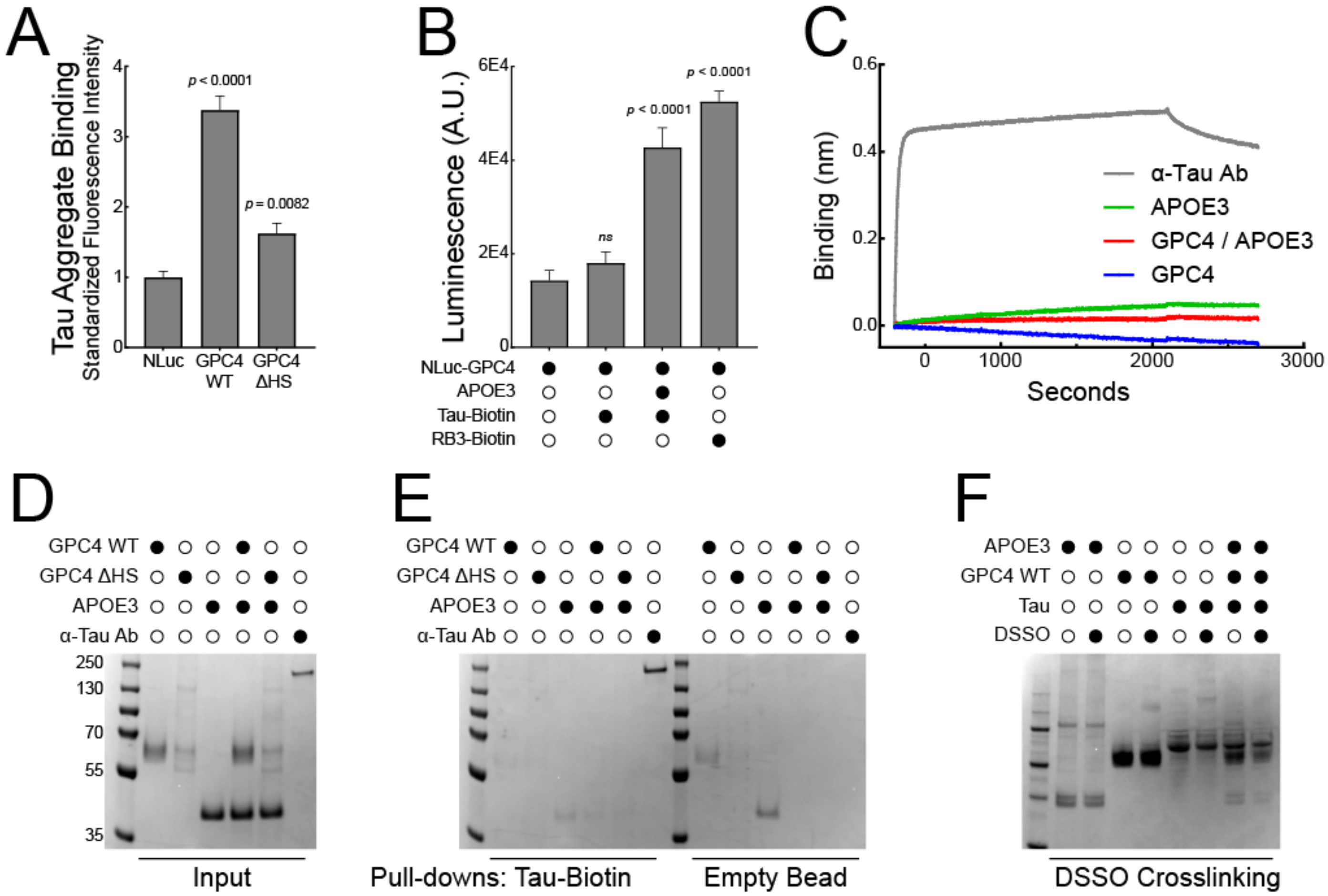


**Figure S8: GPC4, APOE3, and tau interact in a cellular environment.** (A) GPC4 WT, GPC4-ΔHS or a NLuc control plasmids were transiently transfected into HEK293T cells. To measure cell-surface binding, tau-647 fibrils were incubated with the cells for 1 h at 4 °C and cell-surface bound tau was measured via flow cytometry. (B) A constitutively secreted form of NLuc-GPC4 WT was transiently transfected into HEK293T cells and the conditioned media was mixed with monomeric tau-biotin beads with or without APOE3 and pulled down via streptavidin beads. The binding of NLuc-GPC4 WT to tau-biotin was measured via luminescence. The statistical analyses were performed with a one-way ANOVA and Holm-Sidak multiple comparisons tests for the adjusted *p*-values. N = 4 – 8 biological replicates. In all graphs, the data represent the means ± SEM. (C) Biolayer interferometry was used to evaluate direct binding between tau, GPC4, and APOE3. Tau-biotin (100 nM) was immobilized on a streptavidin biosensor before mixing with 250 nM of protein analyte. Only the anti-tau antibody, MD3.1, showed direct binding. (D) SDS-PAGE of recombinant GPC4 WT, GPC4-ΔHS, APOE3, and the anti-tau antibody, MD3.1. (E) Tau-biotin (1 µg) was incubated with 3 µg of the corresponding recombinant purified protein overnight at 4 °C and the complexes were then pulled-down with streptavidin beads, acid eluted, and evaluated via SDS-PAGE. To evaluate non-specific binding, the proteins were co-incubated with streptavidin beads in the absence of tau-biotin (empty bead). Only MD3.1 eluted with the tau-biotin beads. (F) Recombinant purified GPC4 WT, APOE3, and tau were incubated alone or in combination and cross-linked with 50 µM DSSO for 1 h and evaluated via SDS-PAGE. No gel shifts were observed in the presence of DSSO in any condition.
